# Imputation Disparities Driven by Recent Selection and Their Impact on Disease Risk Estimation in East and Southeast Asian Populations

**DOI:** 10.1101/2025.04.03.646928

**Authors:** Dingyang Li, Pattarin Tangtanatakul, Yao Lei, Xiaoxi Liu, Hsi-Yuan Huang, Yang-Chi-Dung Lin, Chengjia Li, Yidan Chen, Lizhi Cai, Jinglu Zhao, Prapaporn Pisitkul, Thanitta Suangtamai, Jinhan Yu, Yihang Zhou, Punna Kunhapan, Rui Sun, Guangjun Yu, Hao Sun, Nattiya Hirankarn, Hsien-Da Huang, Wanling Yang, Yong-Fei Wang

**Affiliations:** School of Medicine, The Chinese University of Hong Kong, Shenzhen, Guangdong, 518172, P.R. China; Warshel Institute for Computational Biology, School of Medicine, The Chinese University of Hong Kong, Shenzhen, Guangdong, 518172, P.R. China; Department of Transfusion Medicine and Clinical Microbiology, Faculty of Allied Health Sciences, Bangkok, 10330, Thailand; Centre of Excellent in Immunology and Immune-Mediated Diseases, Department of Microbiology, Chulalongkorn University, Bangkok, Thailand; Department of Paediatrics & Adolescent Medicine, Queen Mary Hospital, The University of Hong Kong, Hong Kong, P.R. China; Laboratory for Statistical and Translational Genetics, RIKEN Center for Integrative Medical Sciences, Yokohama, Japan; Guangdong Provincial Key Laboratory of Digital Biology and Drug Development, The Chinese University of Hong Kong, Shenzhen, Guangdong, 518172, P.R. China; Faculty of Medicine, Section of Translational Medicine, Mahidol University, Ramathibodi Hospital, Bangkok, Thailand; Division of Allergy, Immunology, and Rheumatology, Department of Medicine, Faculty of Medicine, Ramathibodi Hospital, Mahidol University, Bangkok, Thailand; Department of Medical Sciences, Ministry of Public Health, Nonthaburi, Thailand; School of Medicine, The Second Affiliated Hospital, The Chinese University of Hong Kong, Shenzhen, Guangdong, 518172, P.R. China; Immunology Division, Department of Microbiology, Faculty of Medicine, Chulalongkorn University, Bangkok, Thailand

**Keywords:** Genotype imputation, Whole genome sequencing, GWAS, Polygenic risk score, Population diversity

## Abstract

Using genotype data consisting of 8,316 individuals, we systematically evaluated imputation performance across six state-of-the-art reference panels for Chinese and Thai populations. A substantial proportion of variants identified through whole-genome sequencing, especially low-frequency variants, remained undetected by existing reference panels. In the Chinese population, the TOPMed panel required an R^2^ threshold of 0.60-0.70 to achieve comparable imputation accuracy of the ChinaMAP panel without R^2^ filtering, challenging the standard practice of applying a fixed R^2^ threshold for downstream analyses. Regional analysis highlighted the role of recent selection in imputation discrepancies and revealed an enrichment of immune-related genes in poorly imputed regions. In addition, we showed that the selection of reference panels and R^2^ thresholds could significantly influence estimation of polygenic risk score for disease prediction. These findings underscore the importance of developing ancestrally diverse reference panels and provide valuable guidelines for improving genotype imputation in East and Southeast Asian populations.

## Introduction

Genotype imputation is a statistical method used to infer missing genotypes in individual samples. It is widely applied in genome-wide association studies (GWASs) to increase the density of genetic variants and facilitate meta-analyses across studies with varied genotyping platforms^1^. The accuracy of imputation relies heavily on the characteristics of the reference panel, with sample size, ancestral composition, and haplotype diversity being crucial determinants^2^.

Reference panels produced by the 1,000 Genomes Project^3^ (1KG Phase3 v5; n = 2,504), Haplotype Reference Consortium^4^ (HRC; n = 32,470) and the Trans-Omics for Precision Medicine^5^ (TOPMed r3; n = 133,597) are widely adopted in most GWASs because they offer large sample sizes and rich haplotype data. However, East and Southeast Asian populations remain underrepresented in these reference panels, despite their substantial contribution to global diversity^6,7^. Previous studies have shown that the underrepresentation in the TOPMed reference panel resulted in reduced imputation accuracy for individuals of Asian ancestry compared to those of European ancestry^8^. The selection of appropriate reference panels is essential for uncovering ancestry-specific disease mechanisms and ensuring equitable translational applications of genetic research^9^.

Recent efforts have established specific reference panels for these populations. Notable examples include the Westlake BioBank for Chinese^10^ (WBBC; n = 4,489), China Metabolic Analytics Project^11^ (ChinaMAP; n = 10,155), CHN100k^12^ (n = 25,734) and South and East Asian Reference Database^13^ (SEAD; n = 11,067). However, these reference panels remain underutilized in current studies, partially due to limited evaluation and uncertainty regarding their optimal use in imputation applications.

To enhance the applicability of current reference panels for East and Southeast Asian populations, we curated a SNP array dataset comprising 6,997 individuals of Chinese and Thai ancestry. In addition, we generated a high-coverage whole-genome sequencing (WGS) dataset containing a total of 1,319 individuals from both populations. These datasets enable a comprehensive evaluation of imputation quality across various reference panels.

Imputation quality is commonly evaluated using metrics such as INFO or R^2^, which assess reliability by measuring the proportion of variation explained by the imputation^14^. However, these metrics can become biased when the reference panels inadequately represent the target population, particularly for low-frequency variants^15^. Despite this limitation, most studies apply a fixed threshold (e.g., R^2^ > 0.30) to filter imputed data, potentially compromising downstream analyses.

Genotype concordance rate, which measures the proportion of correctly imputed genotypes for individuals, is widely regarded as the gold standard for evaluating imputation quality. However, this metric can be misleading for variants with skewed allele frequencies. For example, when a variant’s minor allele frequency (MAF) is below 5%, simply predicting the major alleles for all samples can yield over 90% accuracy, regardless of true genotype status^16^. Two alternative metrics address this limitation: the heterozygosity concordance rate, which only evaluates concordance for heterozygous genotypes for individuals and the imputation quality score (IQS)^16,17^, which accounts for chance agreement of genotypes due to biased allele frequency. These metrics provide a more reliable assessment of imputation accuracy.

In this study, we examined imputation quality across six state-of-the-art imputation reference panels for individuals from Chinese and Thai populations and analyzed the underlying factors contributing to imputation disparities. These findings provide valuable guidelines for improving genotype imputation and advancing disease prediction in East and Southeast Asian populations.

## Results

### Dataset preparation

A total of 8,316 samples of Chinese and Thai ancestry were included in this study following quality control (**Methods details; Supplementary Table 1**). The Chinese dataset comprised 3,194 samples previously genotyped using the Infinium Asian Screening Array-24 v1.0 (ASA)^18,19^, a tailored genotyping array for Asian populations, along with 1,263 newly generated WGS samples with an average depth of 36.1. The Thai dataset consisted of 3,803 ASA-genotyped samples from prior studies^18,19^ and 56 newly sequenced WGS samples with an average depth of 43.8.

Genetic variants for the WGS samples were detected using the GATK pipeline^20–22^ (**Methods details**). Variants that passed quality control and overlapped with the ASA platform were extracted from the WGS data. These shared variants were then used to merge the WGS samples with the ASA-genotyped samples. This procedure was performed separately for the Chinese and Thai datasets, resulting in combined ASA-WGS datasets for each population. Principle component analysis (PCA) confirmed no significant difference between the ASA-genotyped and WGS samples within each combined dataset (**Supplementary Figure 1**).

The combined dataset in each population was phased and imputed using six reference panels (**Methods details**): CHN100k, ChinaMAP, WBBC, SEAD panels, and two global reference panels: 1KG (Phase3 v5) and TOPMed (r3). Differences among these reference panels were outlined in **Supplementary Table 2**. Imputation performance was evaluated by comparing the imputed genotypes to the genotypes detected by WGS.

### Coverage of imputation across reference panels

Given that certain reference panels did not support genotype imputation for variants within the human leukocyte antigen (HLA) region (chr6: 29,722,775–33,314,387) or for indels, and showed varied output format for indel imputation, our analysis focused on comparing the proportion of single nucleotide variants (SNVs) outside the HLA region identified through WGS overlapping with those imputed by reference panels. In the Chinese dataset, the imputed variants from CHN100k panel showed the highest overlap with those identified by WGS among the reference panels (68.6%), followed by the TOPMed panel (59.4%) and the ChinaMAP panel (58.9%) (**Supplementary Figure 2a and Supplementary Table 3**). In contrast, the 1KG panel, a widely used global reference, demonstrated the lowest overlap at 31.9%. In the Thai WGS dataset, the SEAD panel exhibited the highest overlap with the WGS-identified variants (81.2%), followed by the CHN100k panel at 78.6%, the TOPMed panel at 77.6% and the ChinaMAP panel at 76.1% (**Supplementary Figure 2b and Supplementary Table 3**). The higher proportion of overlap observed in the Thai dataset may be due to the larger sample size of the Chinese WGS dataset, which enabled the detection of more low-frequency variants.

We further categorized WGS-identified variants as common or low-frequency (MAF < 0.05). As expected, common variants had substantially higher overlap with imputed datasets than low-frequency variants (**Figure 1a and Supplementary Table 3**). In the Chinese dataset, more than 85% of common variants identified by WGS overlapped with those imputed by most reference panels. For low-frequency variants, the CHN100k panel demonstrated the highest overlap at 65.2%, followed by the TOPMed panel at 55.5% and the ChinaMAP panel at 54.3%. Notably, although the 1KG panel showed a high overlap with common variants (89.2%), its overlap for low-frequency variants was considerably lower at 23.6%.

**Figure 1.**
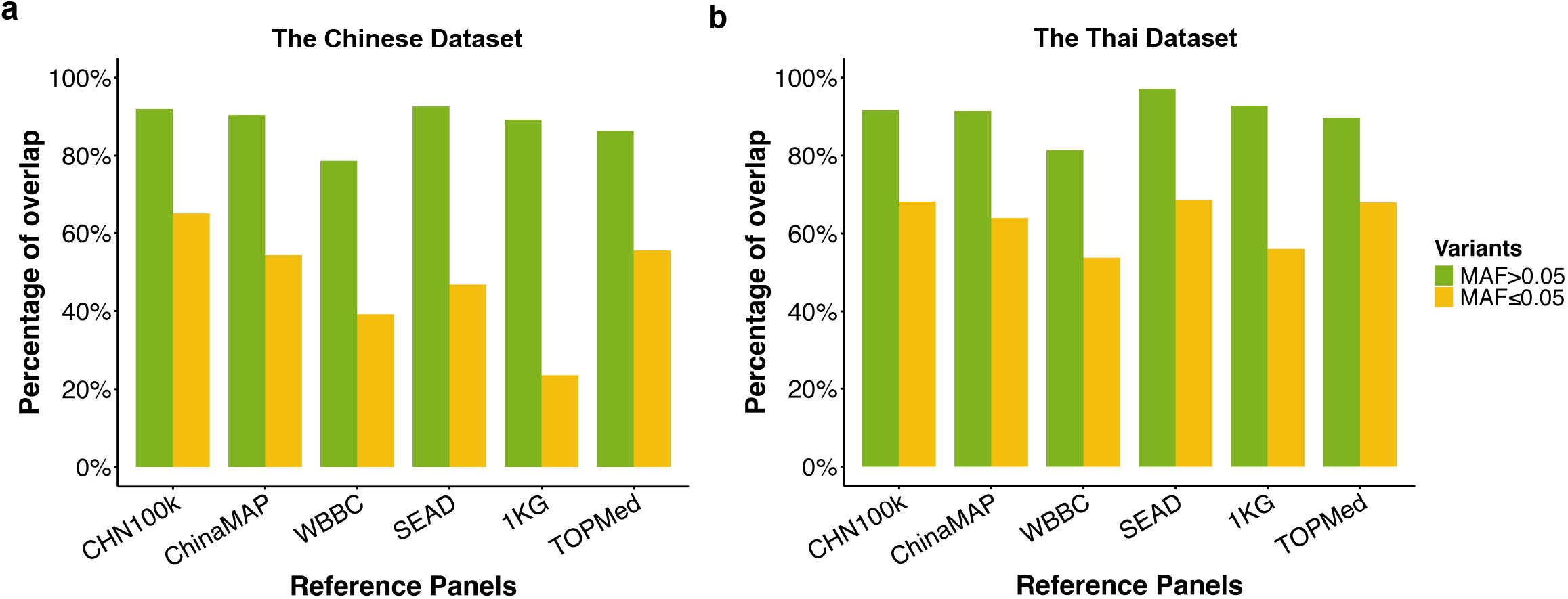
Percentage of WGS-Identified Variants from Chinese (a) and Thai WGS Dataset (b) Overlapping with Imputed Variants Across Reference Panels. The Y-axis represents the percentage of WGS-identified SNVs outside the HLA region that overlap with imputed variants, while the X-axis shows the different reference panels used for genotype imputation. Overlaps for common variants (minor allele frequency > 0.05) are shown in green and overlaps for low-frequency variants (minor allele frequency ≤ 0.05) are displayed in yellow.

A similar pattern was observed in the Thai dataset. 97.1% of common variants identified by WGS overlapped with those imputed by the SEAD panel, followed by the 1KG panel (92.8%), the CHN100k panel (91.6%) and the ChinaMAP panel (91.4%) (**Figure 1b and Supplementary Table 3**). For low-frequency variants, the SEAD panel again demonstrated the highest coverage (68.5%), outperforming the CHN100k (68.1%) and TOPMed (68.0%) panels. In contrast, the 1KG panel showed the low overlap at 56.1%. Collectively, these findings reveal that a significant proportion of WGS-identified variants, particularly low-frequency variants, remain undetected by current imputation references.

### Evaluation of imputation accuracy across reference panels

To evaluate the accuracy of imputed genotypes, the genotypes determined by WGS were considered the true genotypes and used as the benchmark. Variants identified by WGS and shared across all imputation panels were included in the analysis. Accuracy metrics, including imputation R^2^, heterozygosity concordance rate and IQS, were computed for the comparison (**Methods details**). These metrics allowed us to evaluate imputation accuracy at both variant-level (R^2^ and IQS) and sample-level (heterozygosity concordance rate).

In the Chinese dataset, 9,781,349 shared SNVs were included in the analysis. We found that imputation accuracy varied with allele frequency. As expected, common variants generally showed higher imputation accuracy than low-frequency variants across all metrics. Among the reference panels, ChinaMAP consistently outperformed the others, achieving the highest scores across different MAF categories (**Figure 2a**). While the CHN100k panel also performed well, its results were slightly lower than the ChinaMAP. Notably, although the 1KG achieved the second-highest imputation R^2^ scores for common variants, its values appeared inflated relative to actual genotypes. This was evidenced by significantly lower heterozygosity concordance rates and IQS scores compared to ChinaMAP and CHN100k panels (**Figure 2a**).

**Figure 2.**
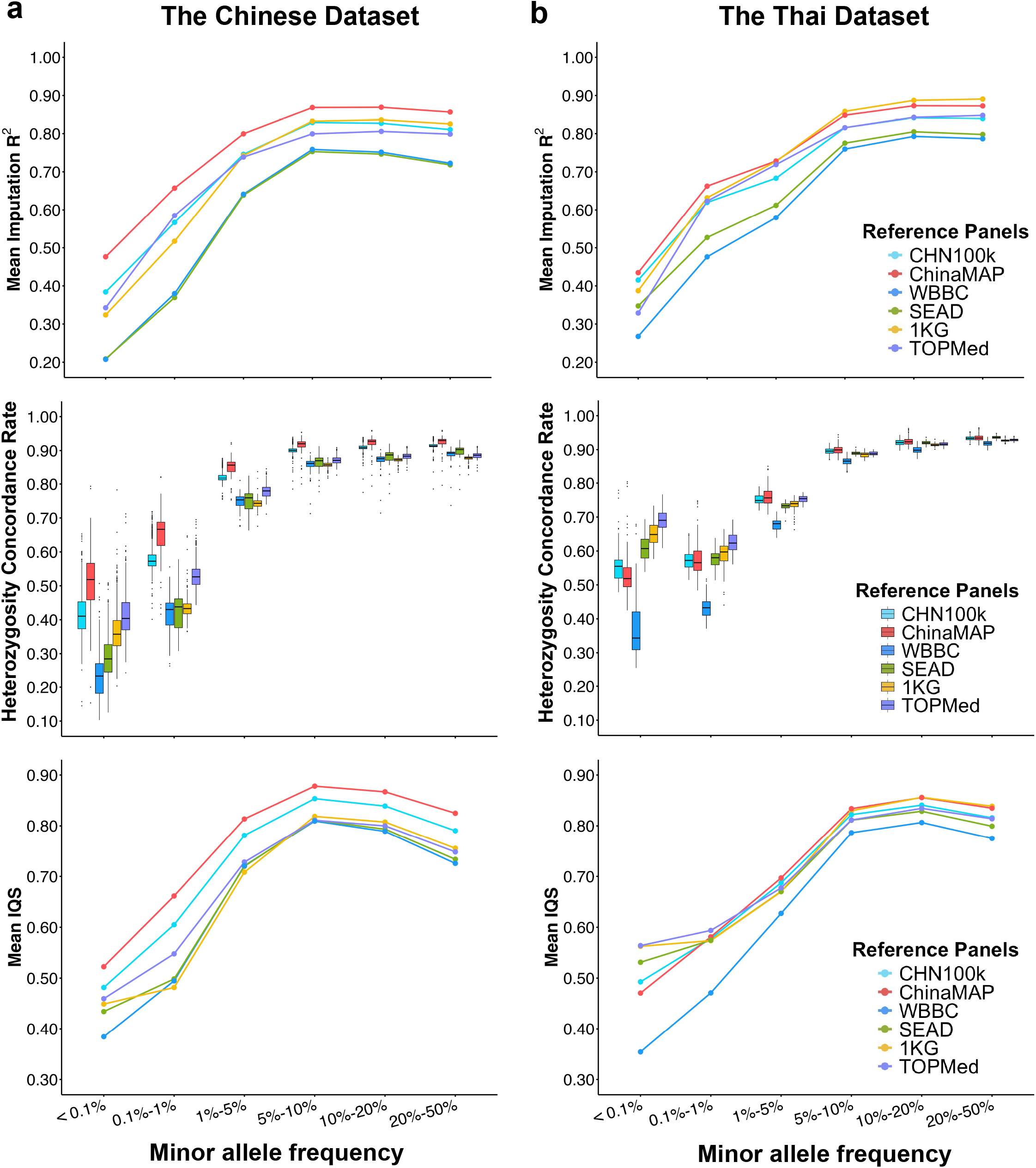
Evaluation of Imputation Quality Across Reference Panels by Minor Allele Frequency Categories in Chinese (a) and Thai (b) WGS Datasets. The X-axis represents minor allele frequency (MAF) categories, while the Y-axis in the top panel displays the mean imputation R2. The middle panel shows the heterozygosity concordance rate for each sample, and the bottom panel represents the mean imputation quality score (IQS) across the MAF categories.

Analysis of the Thai dataset, including 6,299,991 shared variants, revealed different patterns of imputation accuracy across reference panels. Most panels demonstrated comparable performance in both heterozygosity concordance rates and IQS scores across MAF categories, except the WBBC panel (**Figure 2b**). In addition, the TOPMed panel exhibited superior performance for low-frequency variants. Notably, while the SEAD panel demonstrated acceptable concordance rates, we observed consistent deflation in its imputation R^2^ compared to the actual genotypes. Together with the findings in the Chinese dataset, these results suggest that imputation R^2^ scores may not fully reflect the genotype concordance between imputed and actual data among reference panels.

### R^2^ thresholds need to be tailored based on reference panels

Due to the absence of ground-truth genotype data in real-world studies, imputation R^2^ scores remain a critical metric for quality control of imputed variants in GWAS analyses. While current practice commonly employs a fixed R^2^ threshold for this purpose^23^, the above analyses suggest that optimal filtering thresholds should be determined through panel-specific validation studies.

For each reference panel, we used heterozygosity concordance rates and IQS scores as benchmarks to evaluate imputation accuracy across imputation R^2^ thresholds. In the Chinese dataset, while increasing the R^2^ threshold generally improved both heterozygosity concordance rates and IQS scores, the relationships were not strictly linear, particularly for the ChinaMAP and CHN100k panels (**Figure 3a)**. Notably, even without any R^2^ filtering, these two panels achieved a mean heterozygosity concordance rate above 0.90 and an average IQS score exceeding 0.70 (**Figure 3a and Supplementary Table 4-5**). Further increasing the threshold from 0 to 0.40 for the results from the ChinaMAP panel showed minimal improvement, suggesting limited benefit from applying a threshold within this range. By comparison, the 1KG and TOPMed panels required an R^2^ threshold between 0.60 and 0.70 to achieve similar performance levels, which substantially reduced the percentage of imputed variants (**Figure 3a)**.

**Figure 3.**
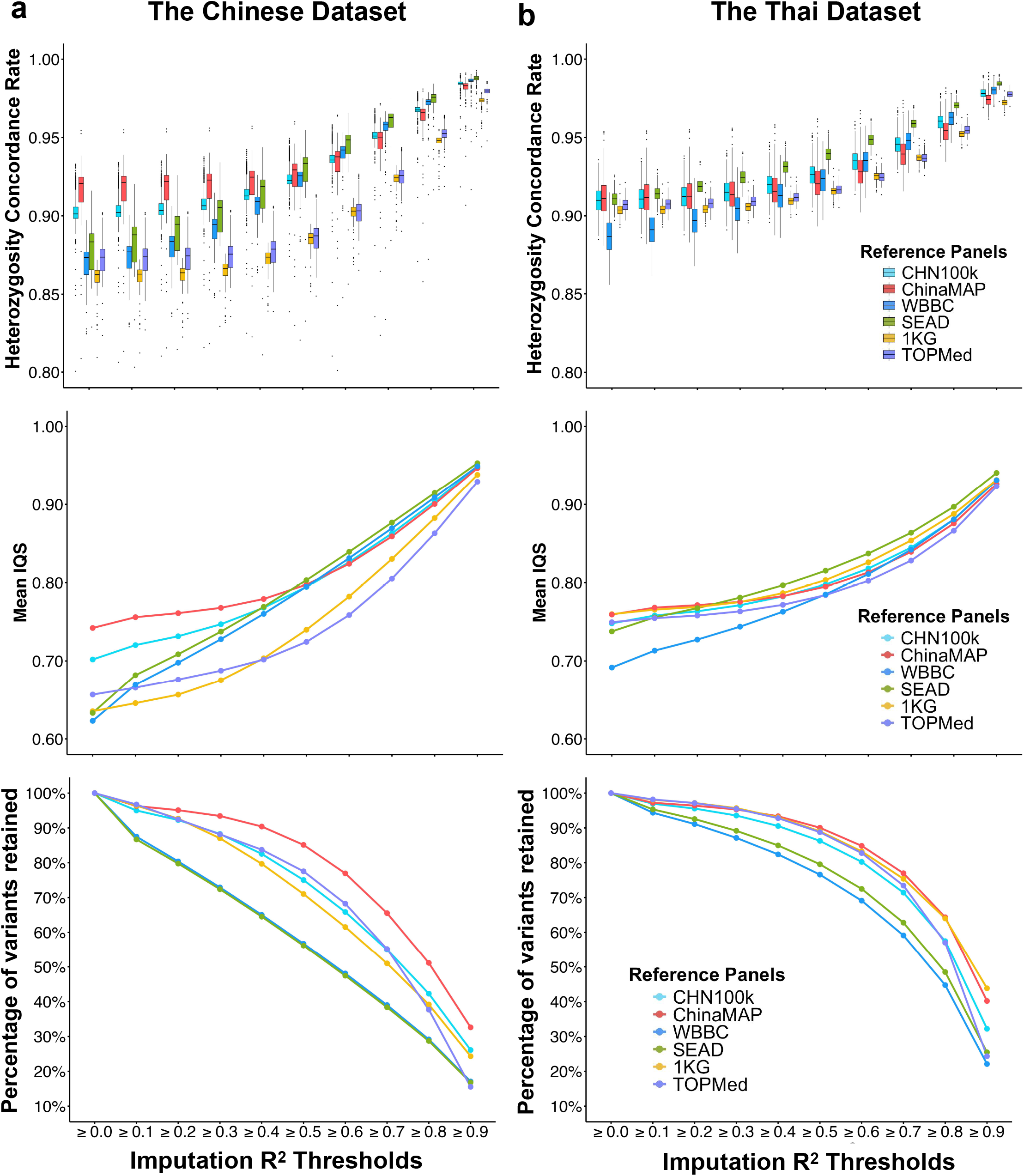
Evaluation of Imputation Quality Across Reference Panels by Imputation R^2^ Thresholds in Chinese (a) and Thai (b) WGS Datasets. The X-axis represents imputation R^2^ threshold categories, while the Y-axis in the top panel displays heterozygosity concordance rate for each sample. The middle panel shows the mean imputation quality score (IQS) and the bottom panel represents percentage of variants retained across R^2^ thresholds.

We also investigated the performance of low-frequency variants. Without R^2^ filtering, the ChinaMAP panel achieved a mean heterozygosity concordance rate above 0.80 and an average IQS score ranging from 0.65 to 0.70 (**Supplementary Figure 3a and Supplementary Table 6-7**). The robust performance underscores the advantage of using the ChinaMAP panel for downstream analyses in Chinese populations.

In the Thai dataset, the SEAD panel slightly outperformed other panels across when the R^2^ threshold was increased to 0.30 (**Figure 3b and Supplementary Table 4-5**). However, without R^2^ filtering, the ChinaMAP and 1KG panels showed considerably higher mean IQS values than the SEAD panel (**Figure 3b**). For low-frequency variants, the heterozygosity concordance rate was substantially lower compared to the Chinese dataset. Achieving a mean heterozygosity concordance rate of 0.80 required raising the R^2^ threshold for the SEAD panel to between 0.40 and 0.50 (**Supplementary Figure 3b and Supplementary Table 6-7**). This adjustment resulted in a reduction of more than 30% of retained low-frequency variants.

Collectively, these findings highlight the critical need to optimize reference panel-specific R^2^ thresholds that account for ancestral background, enabling balanced optimization of imputation accuracy while retaining sufficient variants for downstream analyses.

### Recent selection may contribute to regional variation in imputation quality

We further subdivided autosomal regions into 100 kb windows to assess spatial patterns of imputation quality. Given that allele frequencies vary across genomic regions, we evaluated their performance using the IQS score, which account for biased allele frequency in imputation quality evaluaton^16,17^ (**Methods details**). In the Chinese dataset, the ChinaMAP panel significantly outperformed others, yielding the highest IQS scores in over 92% of windows (**Figure 4a and Supplementary Table 8**). In contrast, in the Thai dataset displayed markedly different patterns: the ChinaMAP and 1KG panels were optimal in approximately 31.7% and 31.1% of windows, respectively, followed by TOPMed (12.7%) and SEAD (11.6%) (**Figure 4b)**. This trend generally aligned with the performance of mean IQS scores observed without R^2^ filtering (**Figure 3b**).

**Figure 4.**
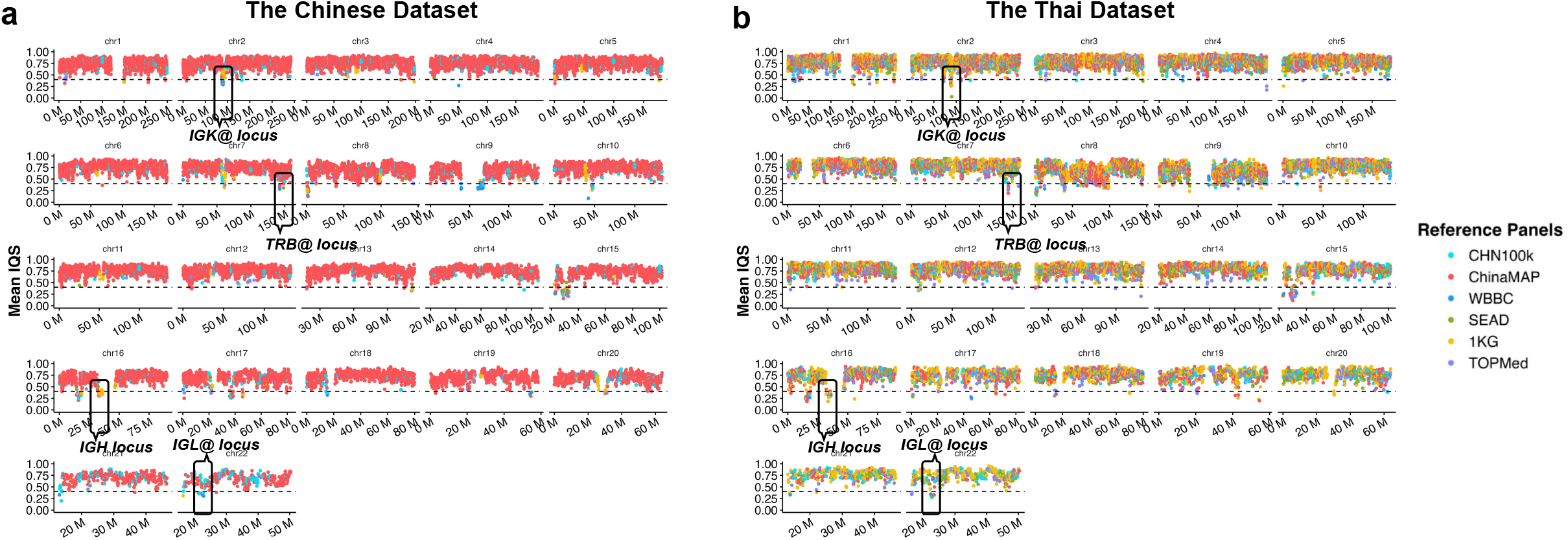
Overview of Reference Panel Performance Across Genomic Regions in the Chinese (a) and Thai (b) Datasets. Each dot corresponds to a 100kb genomic region, with different colors indicating various reference panels. The color of each dot represents the reference panel with the best performance (measured by mean IQS) for that specific region.

We further investigated regional disparities in imputation performance by examining IQS variation among reference panels across genomic windows. In the Chinese dataset, we found 37 windows with substantial differences in imputation quality, characterized by a mean-variance exceeding 0.02 (**Supplementary Figure 4 and Supplementary Table 9)**. Notably, most of these regions (34 out of 37) showed significantly better performance when the Chinese-matched reference panels were used. This observation could be contributed by the effects of recent selection on these regions.

To further evaluate this hypothesis, we examined signatures of recent positive selection using the Singleton Density Score (SDS) method^24^ with data from an independent Chinese cohort^25^ (RePoS database; n = 3,946). Owing to an insufficient number of variants (< 20 variants) within a window, three of the 34 regions were excluded from the analysis. Among the remaining regions, four showed suggestive signatures of recent positive selection (SDS *P*-value < 5E-04). This ratio was significantly higher compared to regions with minimal variation in imputation quality (OR = 6.75, Fisher’s exact test *P*-value = 0.0043; **Method details**).

A notable example is a region on chromosome 11 (55.4 - 55.6 Mb) that contains an *olfactory receptor gene cluster* (**Figure 5a**). Within the region, ChinaMAP (mean IQS = 0.836) demonstrated significantly better imputation performance compared to other panels, such as 1KG (mean IQS = 0.762), TOPMed (mean IQS = 0.644) and SEAD (mean IQS = 0.473). A strong signature of recent positive selection in the region was also detected in the Chinese population, as indicated by the lead SNP rs511492 (SDS = 7.46, SDS *P*-value = 8.69E-14; **Figure 5b**). Prior studies have shown an extended haplotype for the alternative allele (T) compared to the reference allele (C)^25^, further supporting evidence of recent selection in this region. Notably, one of the selected haplotypes (CTT; consisting of rs117749670, rs1459101 and rs511492) carried a stop-gain mutation (rs1459101-C) in the *OR4C16* gene. In our Chinese WGS dataset, this haplotype was observed at a frequency of 15.3%. However, it was nearly absent in African, American, South Asian and European populations (**Figure 5c**). These findings help explain why non-Chinese reference panels performed less effectively in regions that have undergone recent selection in Chinese population.

**Figure 5.**
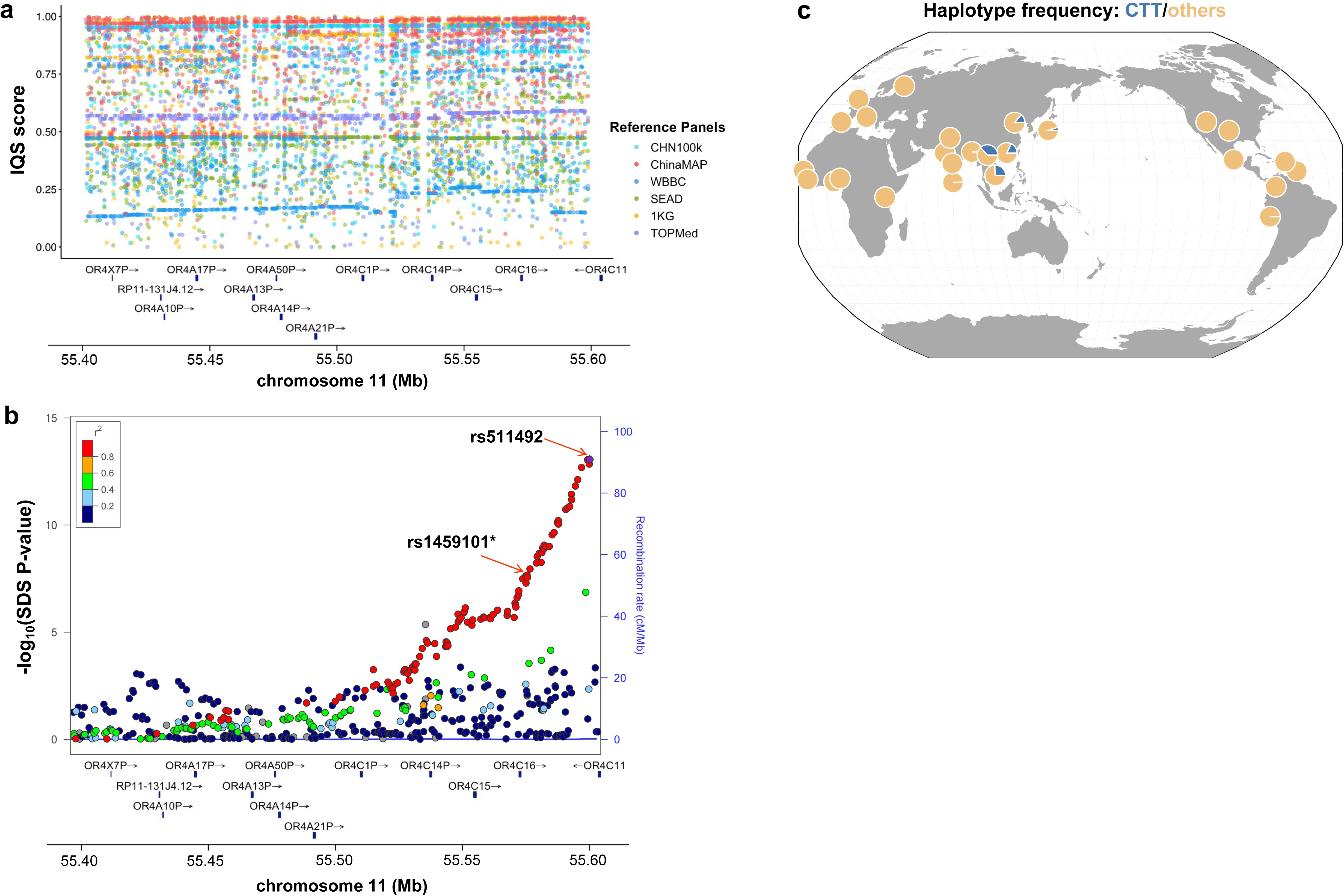
Imputation Disparities and Selection Signatures in the *Olfactory Receptor Gene Cluster* Region. (a) Imputation quality performance across reference panels within the region. Each dot represents individual variants in the region, with different colors indicating the corresponding reference panels. (b) Selection signatures within the region. The recent selection signature is measured by the Singleton Density Score (SDS) method. Each dot represents the –log10 transformed SDS *P*-value for individual variants in the region. *rs1459101 is a stop-gain variant in the *OR4C16* gene. (c) Frequency of the selected haplotype (CTT) across 26 global populations. The CTT haplotype, composed of rs117749670, rs1459101, and rs511492, was analyzed using high-coverage data from the 1,000 Genomes Project (phase 3) to calculate haplotype frequencies.

### Immune-related genes enriched in regions with poor imputation quality

Through the regional evaluation, we also identified 0.94% of genomic regions with consistently poor imputation quality (mean IQS < 0.40) in both populations (**Figure 4 and Supplementary Table 11-12**). These encompassed critical immune-related loci including immunoglobulin heavy loci (*IGH*; 16p11.2), kappa (*IGK@*; 2p11.2) and lambda (*IGL@*; 22q11.22) gene clusters, along with the T cell receptor beta locus (*TRB*; 7q34). These regions pose significant challenges for imputation due to the extensive haplotype diversity, segmental duplications, and complex structural^26–28^. Notably, evidence of recent positive selection at the *IGH* cluster in the Chinese population^25^ underscores the need for long-read sequencing approaches to resolve population-specific architecture in these regions.

Genes within other poorly imputed regions also showed significant enrichment for immunological pathways including macrophage inflammatory protein (*MIP*)-1beta signaling (adjusted *P*-value = 9.20E-06), leukocyte immunoglobulin-like receptor (*LILR*) activity (adjusted *P*-value = 5.02E-05), C-C motif chemokine production (eg. *CCL3, CCL15, CCL18* and *CCL23*; adjusted *P*-value = 1.74E-04) and others (**Supplementary Table 13**). These findings imply that certain association signals for immune-related diseases or traits remain undetected in East and Southeast Asian populations due to inadequate imputation quality.

### Impact of imputation quality on disease risk estimation

Polygenic risk scores (PRS) have been widely utilized to predict individuals’ risk of developing complex diseases^9^. However, the quality of genotype imputation could affect PRS calculations and, consequently, the accuracy of disease prediction. To evaluate this effect, we leverage the recorded phenotypes in Chinese dataset to examine how different imputation references impact PRS estimation to predict individuals’ risk of systemic lupus erythematosus (SLE).

Using WGS from 1,263 Chinese individuals, we compared SLE PRS calculated from true genotypes against the PRS generated by genotype dosages imputed from each reference panel (**Methods details**). To compute the PRS, we used 203 previously reported SLE-associated variants^19,29^ shared between the WGS dataset and the reference panels. The results showed that PRSs calculated from imputation using the ChinaMAP (r = 0.945) and CHN100K (r = 0.937) panels demonstrated much stronger correlations with the score derived from the true genotypes, compared to PRSs computed using other reference panels (e.g., r = 0.894 for 1KG, r = 0.898 for TopMed; **Figure 6a**).

**Figure 6.**
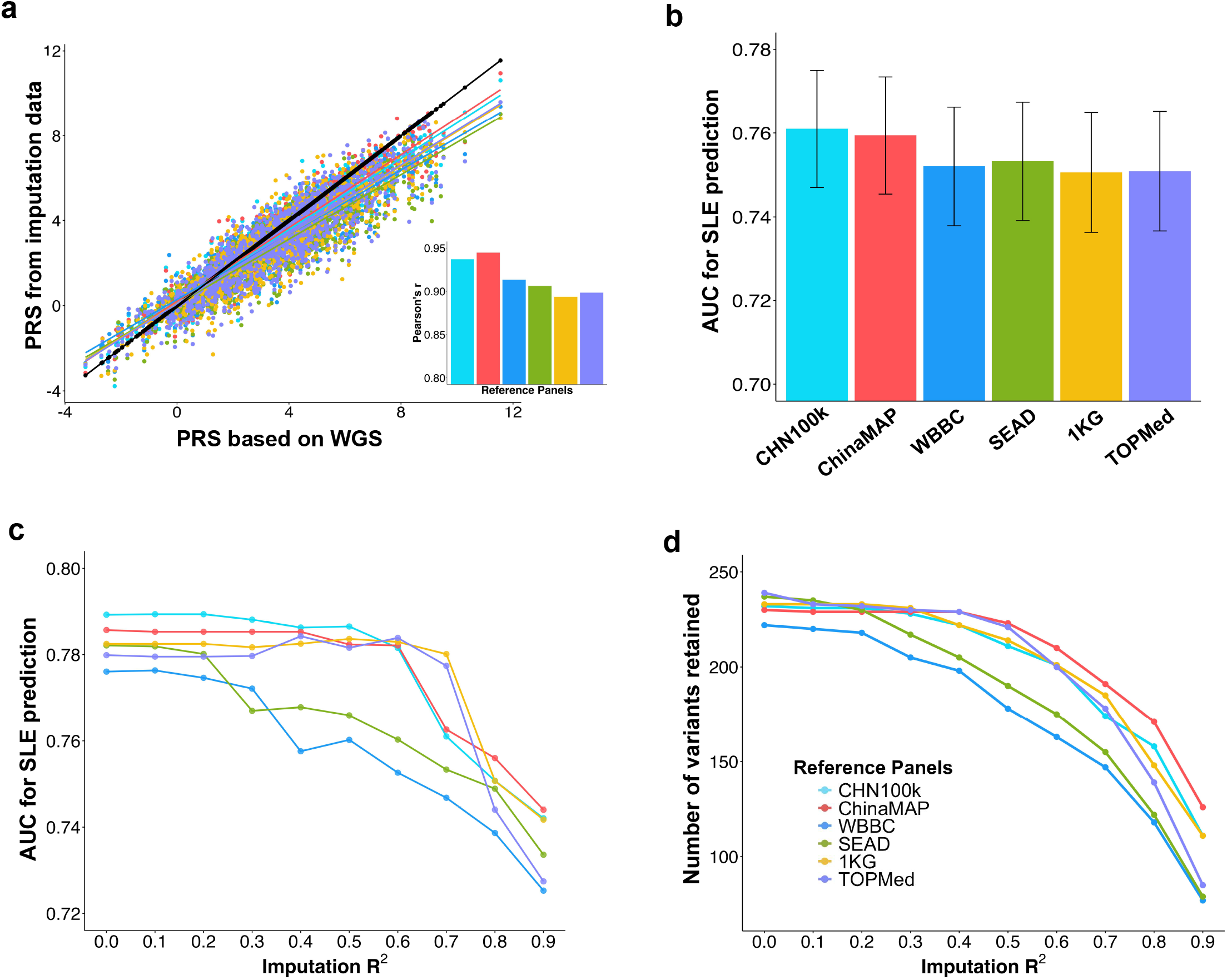
Impact of Imputation Results on Disease Risk Prediction. (a) Comparison of polygenic risk scores (PRS) derived from true genotypes with PRS computed using genotype dosages imputed from various reference panels. Each dot represents an individual WGS sample. (b) Predictive accuracy for systemic lupus erythematosus (SLE) using PRS computed by various reference panels. The accuracy is measured by the area under the receiver operating characteristic curve (AUC). Error bars represent the 95% credible intervals for AUC values. (c) Predictive accuracy for SLE using PRS built from variants filtered at different R^2^ thresholds. (d) The number of SLE-associated variants included in constructing the PRS at various R^2^ thresholds.

To further evaluate the impact on disease risk prediction, we extended the analysis to the ASA-WGS combined dataset, comprising 2,027 SLE cases and 2,430 controls of Chinese ancestry. Using this data, we measured the accuracy of SLE prediction by the area under the receiver-operating characteristic curve (AUC). The results showed that the PRSs constructed using genotype dosage imputed from the CHN100k (AUC = 0.761) and ChinaMAP (AUC = 0.759) panels performed slightly better in SLE prediction than those generated using other panels (e.g., AUC = 0.751 for both 1KG and TopMed; **Figure 6b**).

We next investigated the impact of varying R^2^ thresholds on prediction performance. Unlike the above study, all 239 previously reported SLE-associated variants (**Supplementary Table 15**) were included in this analysis. The results showed that overall predictive power declined as the R^2^ threshold increased (**Figure 6c**). The mean AUC across panels dropped significantly from 0.789 at an R^2^ threshold of 0 to the mean of 0.725 at an R^2^ threshold of 0.90 (Paired t-test *P*-value = 2.31E-06). This reduction is likely attributed to the decreased number of associated variants retained at higher R^2^ thresholds (**Figure 6d)**.

In addition, we observed that PRSs derived from the CHN100k and ChinaMAP panels achieved the best performance without any R^2^ filtering. In contrast PRSs from the TOPMed and1KG panels performed optimally at an R^2^ threshold of 0.40 (**Figure 6c**). This pattern suggests that raising the threshold may enhance predictive power by reducing the influence of poorly imputed genotypes when utilizing global reference panels. Collectively, these findings highlight the importance of ancestry-matched reference panels in improving PRS estimation and disease risk prediction while providing insights into optimizing thresholds for global panels.

## Discussion

The underrepresentation of East and Southeast Asian populations in widely used global reference panels, such as TOPMed, poses significant challenges for genetic discovery and translational applications in these ancestral groups. By leveraging high-coverage WGS data as a gold standard, we sought to address longstanding uncertainties regarding the optimal selection of imputation panels and quality control strategies for GWAS in East and Southeast Asian populations.

Our study suggests that a substantial proportion of WGS-identified variants, particularly low-frequency variants, remain uncovered by current imputation references. While global reference panels effectively capture common variants, ancestry-matched panels provide significantly better coverage for low-frequency variants. Notably, we showed an often-overlooked limitation of earlier studies that relied on the 1KG panel, which showed the lowest overlap for low-frequency variants in East and Southeast Asian populations. These results emphasize the continued importance of sequencing technologies for comprehensive variant discovery in future studies.

Our findings demonstrate that imputation R^2^ scores do not accurately represent the true concordance between imputed and actual genotype data, and the concordances at a fixed R^2^ threshold vary considerably across different reference panels. For example, in our analysis of Chinese data, the ChinaMAP panel achieved a good performance without any R^2^ filtering. In contrast, the TOPMed panel required an R^2^ threshold between 0.60 and 0.70 to reach comparable performance. The deviation may result from genetic differences between the study and reference populations^30^. Therefore, panel-specific R^2^ thresholds should be considered rather than applying a uniform threshold for quality control.

This study provides actionable guidelines for genotype imputation and quality control in GWAS of East and Southeast Asian populations. For studies involving Chinese population, the ChinaMAP and CHN100k panels are recommended due to their superior coverage and accuracy compared to other reference panels. ChinaMAP showed the best performance in imputation accuracy. However, this panel does not support the imputation of indels and SNVs within the HLA region. Given the high genetic diversity of HLA alleles and their critical role in immune response and disease susceptibility, there is a need for specialized tools tailored to high-quality HLA allele imputation.

For Thai population studies, the SEAD panel would be the preferred choice, as it demonstrated broader coverage and slightly higher concordance with actual genotypes when the R^2^ threshold is increased to 0.30. However, regional analysis revealed that this panel achieved the best performance in only 11.6% of genomic regions, significantly lower than the performance observed for the ChinaMAP and 1KG panels. This finding suggests that researchers may need to select reference panels based on specific genomic regions of interest when ancestry-matched panels are unavailable. In addition, the Thai dataset exhibited greater variation in imputation quality, with 168 genomic windows showing significant differences across reference panels, compared to 37 windows in the Chinese dataset (**Supplementary Figure 4 and Supplementary Table 10**). These findings highlight the higher heterogeneity in imputation quality for Thai populations and underscore the need for more tailored and optimized reference panels to enhance imputation accuracy for this population.

More importantly, our study further suggests that recent selection may contribute to the imputation variations across different reference panels. This is exemplified by the olfactory receptor gene cluster on chromosome 11, which has been reported to be under recent positive selection in the Chinese population^25^ (**Figure 5**). The selected haplotype (CTT) in this region is highly prevalent in East Asian populations but nearly absent in others. As the unique haplotype structure, imputation quality can be significantly improved when using ancestry-matched reference panels. Previous studies on the lactase (*LCT*) gene also highlight potential limitations of genotype imputation in the regions under recent selections^31^. In addition, recent investigations in type 2 diabetes (T2D)-associated loci demonstrated that haplotype differences between study cohorts and the reference panels can skew imputation results, favoring allelic calls that are more common in the reference panel^32^. Taken together, these findings underscore the importance of using a matched reference panel to improve imputation accuracy in regions with varying haplotype frequencies across populations.

Our studies also revealed that certain regions, including multiple immune-related genes such as the *IGH, IGK@, IGL@* and *TRB* loci, were poorly imputed in East and Southeast Asian Populations. The complex structure and evidence of recent selection within these regions^26–28^ suggest that targeted long-read sequencing approaches are necessary to refine them in an ancestry-specific manner. In addition, genes involved in *MIP*-1beta signaling, *LILR* activity and C-C motif chemokine production, playing an important role in the immune system^33–36^, were also enriched in these poorly imputed regions. These findings suggest that certain essential association signals related to immune-related diseases or traits may be ignored in previous studies. However, it is important to note that these results may be influenced by the SNP array platform used in our analysis, which was limited to the Infinium ASA platform. Further studies are needed to determine whether imputation quality in these regions can be improved using other genotyping platforms.

Finally, we demonstrated the significant impact of reference panel selection on disease risk prediction, using SLE as a model disease. Our results showed notable improvements in predictive power when using ancestry-optimized reference panels. Notably, filtering variants based on an R^2^ threshold would reduce predictive power when using an ancestry-matched reference panel. In contrast, we recommend raising the R^2^ threshold when relying on a global reference panel to enhance prediction power. However, since our analysis focused solely on SLE, further studies are needed to determine whether these patterns hold for other diseases.

In summary, this study provides valuable guidelines for future GWAS in East and Southeast Asian populations. It underscores the importance of continued efforts to develop ancestrally diverse reference panels. Together with advancements in long-read sequencing technologies, these efforts will be essential for addressing current disparities in imputation quality and enhancing the accuracy of disease prediction.

## STAR★Methods

### Key resources table

| REAGENT or RESOURCE | SOURCE | IDENTIFIER |
| --- | --- | --- |
| **Deposited data** |  |  |
| 1000 Genomes Project Phase 3 | The 1000 Genomes Project Consortium.<sup>3</sup> | https://www.internationalgenome.org/ |
| Genetic map (hg38) | The International HapMap Consortium.<sup>37</sup> | https://storage.googleapis.com/broad-alkesgroup-public/Eagle/downloads/tables/genetic_map_hg38_withX.txt.gz |
| RePoS SDS dataset | Luo et al.<sup>25</sup> | http://bigdata.ibp.ac.cn/RePoS/data/download/ |
| **Software and algorithms** |  |  |
| CHN100k Imputation Server | Jiang et al.<sup>12</sup> | http://bioinformatics.hit.edu.cn/imputation/ |
| ChinaMAP Imputation Server | Li et al.<sup>38</sup> | http://www.mbiobank.com/imputation/ |
| Westlake Imputation Server | Cong et al.<sup>10</sup> | https://imputationserver.westlake.edu.cn/index.html |
| TOPMed Imputation Server | Taliun et al.<sup>5</sup> | https://imputation.biodatacatalyst.nhlbi.nih.gov/ |
| Michigan Imputation Server | Das et al.<sup>14</sup> | https://imputationserver.sph.umich.edu/index.html |
| PLINKv1.9 | Chang et al.<sup>39</sup> | http://www.cog-genomics.org/plink/1.9/ |
| EAGLE2 | Loh et al.<sup>40</sup> | https://alkesgroup.broadinstitute.org/Eagle/ |
| VCFtools | Danecek et al.<sup>41</sup> | https://vcftools.sourceforge.net/ |
| GLnexus | Michael F et al.<sup>42</sup> | https://github.com/dnanexus-rnd/GLnexus |
| Clara Parabricks (v4.3.0-1) | NVIDIA (2024).<sup>43</sup> | https://www.nvidia.com/en-us/clara/genomics/ |
| Toppgene | Chen et al.<sup>44</sup> | https://toppgene.cchmc.org/ |
| Locuszoom | Boughton et al.<sup>45</sup> | https://github.com/statgen/locuszoom |
| pROC package | Robin et al.<sup>46</sup> | https://www.expasy.org/resources/proc |

## Resource availability

### Lead contact

The lead contact for this paper is Yong-Fei Wang

### Materials availability

This study did not generate new reagents.

### Data and code availability

The details of tools and public datasets used for conducting this research are provided in the key resource table. Code for variant calling, quality control, and IQS calculation is publicly available at: https://github.com/LI-DINGYANG/Synecdoche. Any additional information required to reanalyze the data reported in this paper is available from the lead contact upon request.

## Methods details

### Sample overview

This study collected a total of 8,438 individuals from Chinese and Thai populations. Of these, 3,198 Chinese samples and 3,830 Thai samples were sourced from our earlier studies and genotyped using the Infinium Asian Screening Array-24 v1.0 (ASA)^18,19^. To assess imputation accuracy, we recruited 1,353 samples of Chinese ancestry from Hong Kong, China and 57 samples of Thai ancestry from Bangkok, Thailand. Whole-genome DNA from these individuals was sequenced using the DNBSEQ platform by BGI Genomics, with a read length of 150 base pairs (PE150).

### Variant calling

NVIDIA Parabricks v4.3.0 software suite^43^ was used to identify germline variants for WGS data. The software suite significantly accelerated genomic analysis by utilizing graphics processing units (GPUs) instead of traditional central processing units (CPUs), achieving faster processing speeds while maintaining accuracy equivalent to the standard GATK (Genome Analysis Toolkit) best practices^20–22^.

The analysis followed a series of steps to process the sequencing data. Raw paired-end sequencing data were aligned to the GRCh38 reference genome using the BWA-mem algorithm. PCR-induced duplicate fragments were removed with the MarkDuplicates algorithm. Base Quality Score Recalibration (BQSR) was applied to correct sequencing quality discrepancies that arise from different sequencing cycles and contexts. Variants in individual samples were subsequently detected using the HaplotypeCaller algorithm. Joint genotyping was performed using Glnexus^42^, a tool designed for large-scale genomic variant analysis.

In addition, vcftools^41^ was used to refine the data by filtering out variants with genotype quality (GQ) less than 20 or depth less than 5. Variants with missing rate exceeding 20% among samples were removed. “Half-called” variants, referring to genotypic calls that are incomplete or ambiguous within the variant calling process, were excluded from the analysis. Following these steps, 46,218,832 variants on autosomes were retained in the Chinese WGS dataset, consisting of 42,751,494 SNVs and 3,467,338 indels. Similarly, 15,604,227 variants on autosomes were retained in the Thai WGS dataset, including 12,598,039 SNVs and 3,006,188 indels. The transition/transversion (Ti/Tv) ratio for individuals ranged from 2.03 to 2.06 in both datasets.

### Quality control and genotype imputation

To enhance phasing accuracy, the WGS samples were merged with the ASA-genotyped samples based on their shared variants. Quality control steps were applied to the merged datasets using PLINKv1.9^39^. Variants with missingness greater than 0.05, MAF less than 0.01, or failing the Hardy-Weinberg equilibrium (HWE) test (*P*-value < 1.00E-05) were removed. Samples identified as potential duplicates (identical-by-descent relationship PIHAT > 0.90), or exhibiting abnormal heterozygosity levels (|F-score| > 0.1) were also excluded. After quality control, 8,316 samples from both populations were retained for downstream analyses (**Supplementary Table 1**).

Eagle (version 2.4.1)^40^ was then used to phase samples from the WGS-ASA combined dataset for each population without using reference. The Genetic map generated by HapMap project^37^ was utilized during this process. The phased datasets from Chinese and Thai populations were submitted to the CHN100k^12^, ChinaMAP^38^, Westlake^10^, TOPMed^5^ and Michigan^14^ Imputation Servers, respectively. The 1KG reference panel was included in the Michigan Imputation Server and the SEAD reference panel was included in the Westlake Imputation Server. The Minimac4 algorithm^14^ was used for genotype imputation across all imputation servers.

### Evaluation of imputation quality

Heterozygosity concordance rate and IQS score^16,17^ were used to evaluate imputation performance across reference panels. The heterozygosity concordance rate was calculated by comparing the observed heterozygous genotypes from WGS with the imputed genotypes for each individual. The IQS score for each variant was computed by subtracting the chance agreement (*P*_*c*_) from the observed agreement (*P*_*o*_), and then dividing the result by the maximum possible agreement excluding the chance agreement (Equation 1). The calculation is represented by the following equations^16,17^:

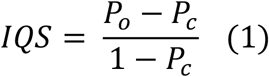

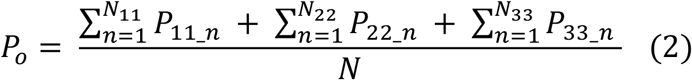

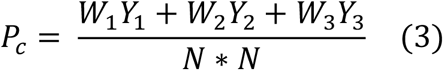

Where *P*_*o*_ represents the observed concordance rate for different genotype classes, calculated as the sum of estimated genotype probabilities for each matching genotype class (Equation 2). *P*_11_ denotes the estimated genotype probability for the homozygous reference genotype (0/0) in cases where the true genotype, determined by WGS, is 0/0 and the imputed genotype also matches as 0/0. Similarly, *P*_22_ and *P*_33_ represent the estimated genotype probabilities for the heterozygous genotype (0/1) and the alternative homozygous genotype (1/1), respectively, in cases where both the true and imputed genotypes align. *N*_11_, *N*_22_ *and N*_33_ refer to the number of individuals with matched genotypes for the homozygous reference (0/0), heterozygous (0/1) and alternative homozygous (1/1) categories, respectively. *N* denotes the total number of individuals.

The chance agreement (*P*_*c*_) is calculated as the sum of the products of marginal frequencies, representing the expected agreement if genotypes were randomly assigned based on their respective marginal rates (Equation 3). Specifically, *W*_*1*_, *W*_*2*_ and *W*_*3*_ refer to the marginal frequency for the observed genotype 0/0, 0/1 and 1/1, respectively, while *Y*_*1*_, *Y*_*2*_ and *Y*_*3*_ represent the marginal frequency for the imputed genotype 0/0, 0/1 and 1/1, respectively, as illustrated in **Supplementary Table 14**.

For the analysis of the Chinese dataset, the MAF categories were defined using 1,263 Chinese WGS samples. For the analysis of the Thai dataset, the MAF categories were defined based on WGS data derived from 1KG East Asian populations, due to a limited number of WGS samples in the Thai dataset.

### Region-based analysis

For regional analysis, 9,781,349 SNVs shared by all reference panels and the Chinese WGS dataset were used in the Chinese dataset. Similarly, 6,299,991 shared SNVs were used for analysis in the Thai dataset. To conduct the analysis, the genome was divided into windows of 100 kbp. This resulted in 26,352 windows in the Chinese dataset and 26,029 windows in the Thai dataset, excluding the HLA region. The difference in the number of windows between the two datasets reflected different variants included in each analysis (**Supplementary Table 8**).

To account for differences in allele frequencies across genomic regions, we assessed imputation quality using the IQS score. For each genomic window, the mean IQS score for each reference panel and the variation in IQS across panels were calculated. Windows with a mean IQS below 0.4 across all panels were classified as regions with poor imputation quality, while those with a variance in IQS exceeding 0.02 across panels were categorized as regions with significant differences. Regions identified as having poor imputation quality in both the Chinese and Thai datasets were further analyzed. From these regions, 542 genes were identified and used for enrichment analysis with ToppGene^44^ (**Supplementary Table 13**).

To investigate whether genomic regions with substantial variation in imputation accuracy across reference panels were enriched in regions under recent positive selection, we analyzed selection signals using the SDS method^24^. The SDS data were sourced from the RePoS database, including 3,946 Chinese individuals^25^. We focused on regions with the best performance in Chinese ancestry-matched panels. We excluded regions with an insufficient number of variants (< 20 variants) in the RePoS database. For comparison, a set of 22,540 genomic regions with an IQS variance of less than 0.005 was used as a benchmark. Fisher’s exact test was then conducted to examine whether genomic regions with substantial variation in imputation accuracy were more likely to be enriched in regions under recent positive selection compared to regions without significant variation. Due to insufficient data for assessing recent positive selection in the Thai population, this analysis was not performed on the Thai dataset.

### Calculation of polygenic risk scores

The polygenic risk score (PRS) of SLE was calculated for each individual using the equation provided below:

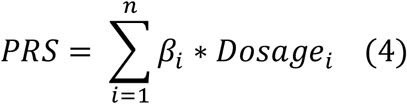

Where *β*_*i*_ represents the effect size of the i_th_ SLE-associated variant, *Dosagei* represents the corresponding allele dosage obtained from imputation results, and n is the total number of associated variants used in the calculation. In this study, a total of 239 SLE-associated variants outside of the HLA region were extracted from previous studies^19,29^ and their genetic effect on SLE development was summarized in **Supplementary Table 15**.

To assess the impact of reference panel selection on SLE prediction, the PRS was calculated using 203 SLE-associated variants that were shared across the six reference panels and the Chinese WGS datasets. No R^2^ filtering was applied in this analysis. For the WGS samples in the Chinese population, we evaluated the accuracy of PRSs by comparing those derived from imputed genotype dosages with PRSs calculated from sequencing-based genotypes, using the Pearson Correlation Coefficient (r) to measure the agreement.

The performance of various reference panels on SLE prediction for the ASA-WGS combined dataset was assessed using the AUC score. To examine the impact of different imputation R^2^ thresholds on SLE prediction, all SLE-associated variants were analyzed. For each R^2^ threshold, PRSs were calculated using the associated variants retained in each reference panel, and the corresponding AUC scores were calculated. The AUC scores and the 95% confidence intervals were calculated using the pROC^46^ package.

## Ethics approval

This study was approved by the institutional review boards, including the ethical committee from the Hospital Authority Hong Kong West Cluster (UW 07-119) and the Faculty of Medicine Ramathibodi Hospital, Mahidol University (12-58-12).

## Acknowledgements

We gratefully acknowledge funding support from the Shenzhen-Hong Kong Jointly Funded Project (Category A; SGDX20230116093201002) and the Stability Support for Higher Education from Shenzhen Science and Technology Program, the Guangdong Natural Science Foundation Youth Enhancement Project (2024A1515030287), the Guangdong S&T programme (2024A0505050001), and the National Natural Science Foundation of China (Grant No.82471825). We also extend our thanks to the Warshel Institute for Computational Biology and their funding support from Shenzhen City and Longgang District (LGKCSDPT2024001). The authors thank the anonymous reviewers for their constructive comments and valuable suggestions that helped to improve the quality of the manuscript.

## Author contributions

Y.-F. Wang conceived the study. D. Li took the lead in data analysis. P. Tangtanatakul, Y. Lei, H-Y. Huang, Y.-C.-D. Lin, C. Li, Y. Chen, L. Cai, J. Zhao, P. Pisitkul, T. Suangtamai, J. Yu, Y. Zhou, P. Kunhapan, R. Sun, G. Yu, H. Sun, N. Hirankarn, H.-D, Huang, W. Yang undertook subject recruitment and collected phenotype data. D. Li, P. Tangtanatakul, Y. Lei, X. Liu, W. Yang and Y.-F. Wang carried out data analyses and interpretation. D. Li, P. Tangtanatakul and Y.-F. Wang wrote the manuscript. All authors read and contributed to the manuscript.

## Inclusion and diversity

We support inclusive, diverse, and equitable conduct of research.

