## Supplemental Figures for "Imputation Disparities Driven by Recent Selection and Their Impact on Disease Risk Estimation in East and Southeast Asian Populations"

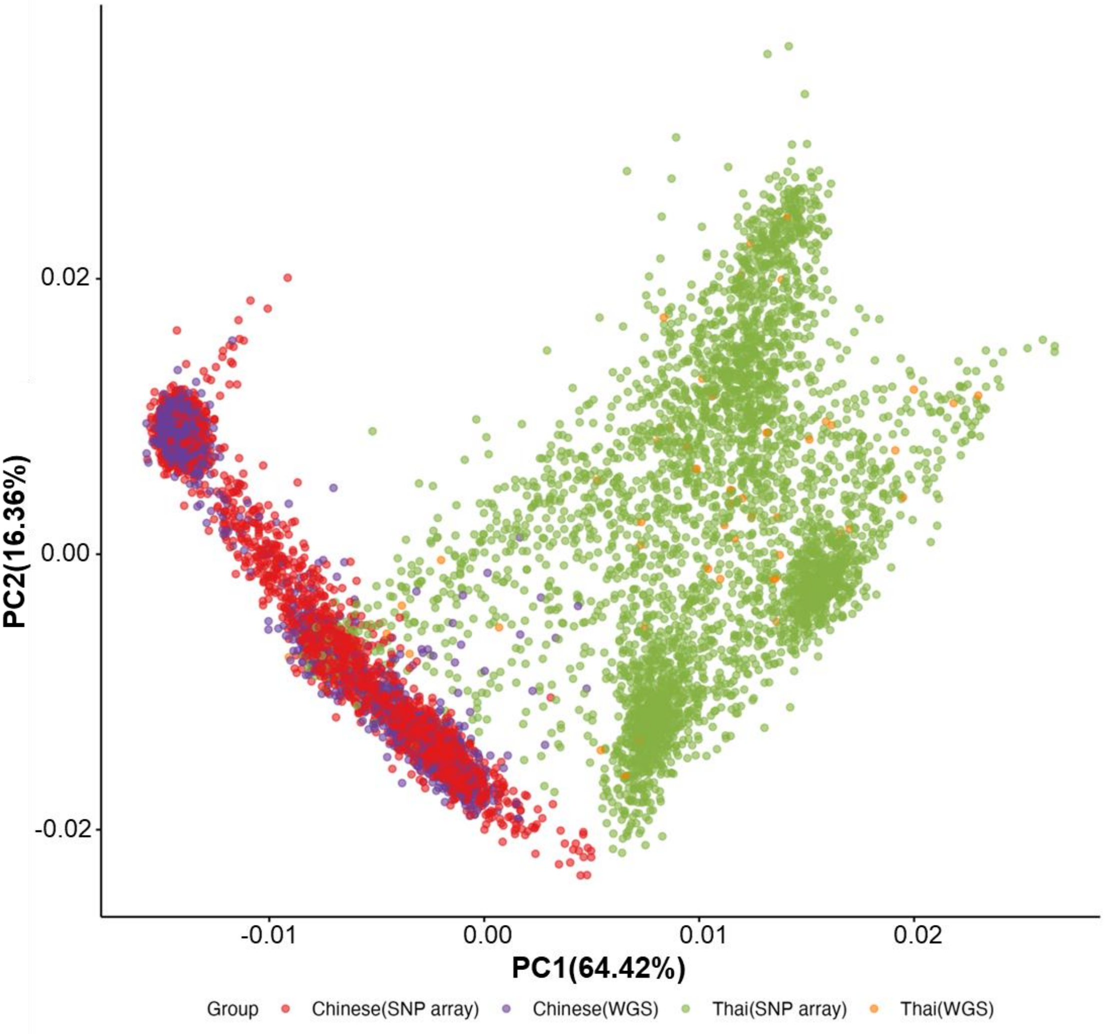


**Supplementary Figure 1 Principal Component Analysis (PCA) of Samples from Whole Genome Sequencing (WGS) and SNP Arrays in the Chinese and Thai Datasets**

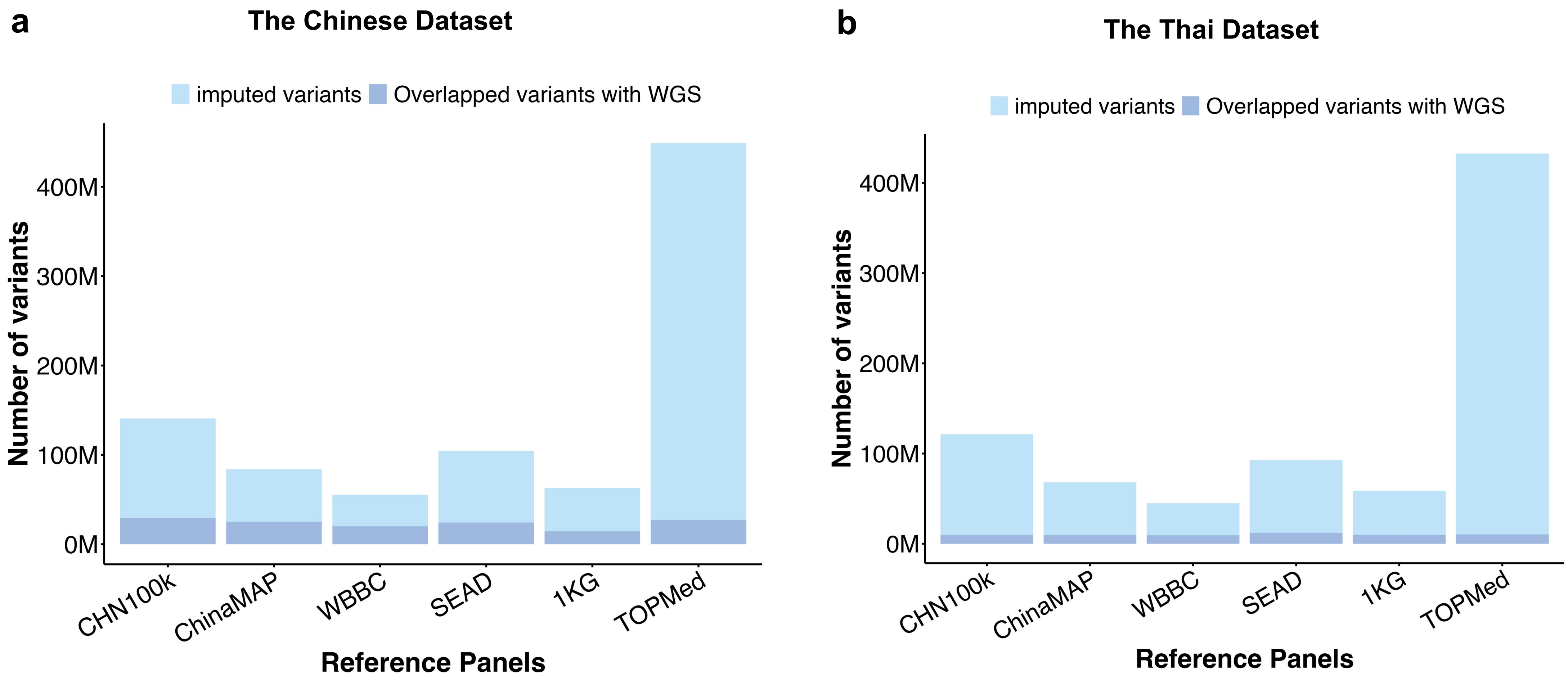


**Supplementary Figure 2 Summary of Imputed Variants Across Reference Panels and Their Overlap with Variants Detected by Whole Genome Sequencing (WGS) in Chinese (a) and Thai Datasets (b)**

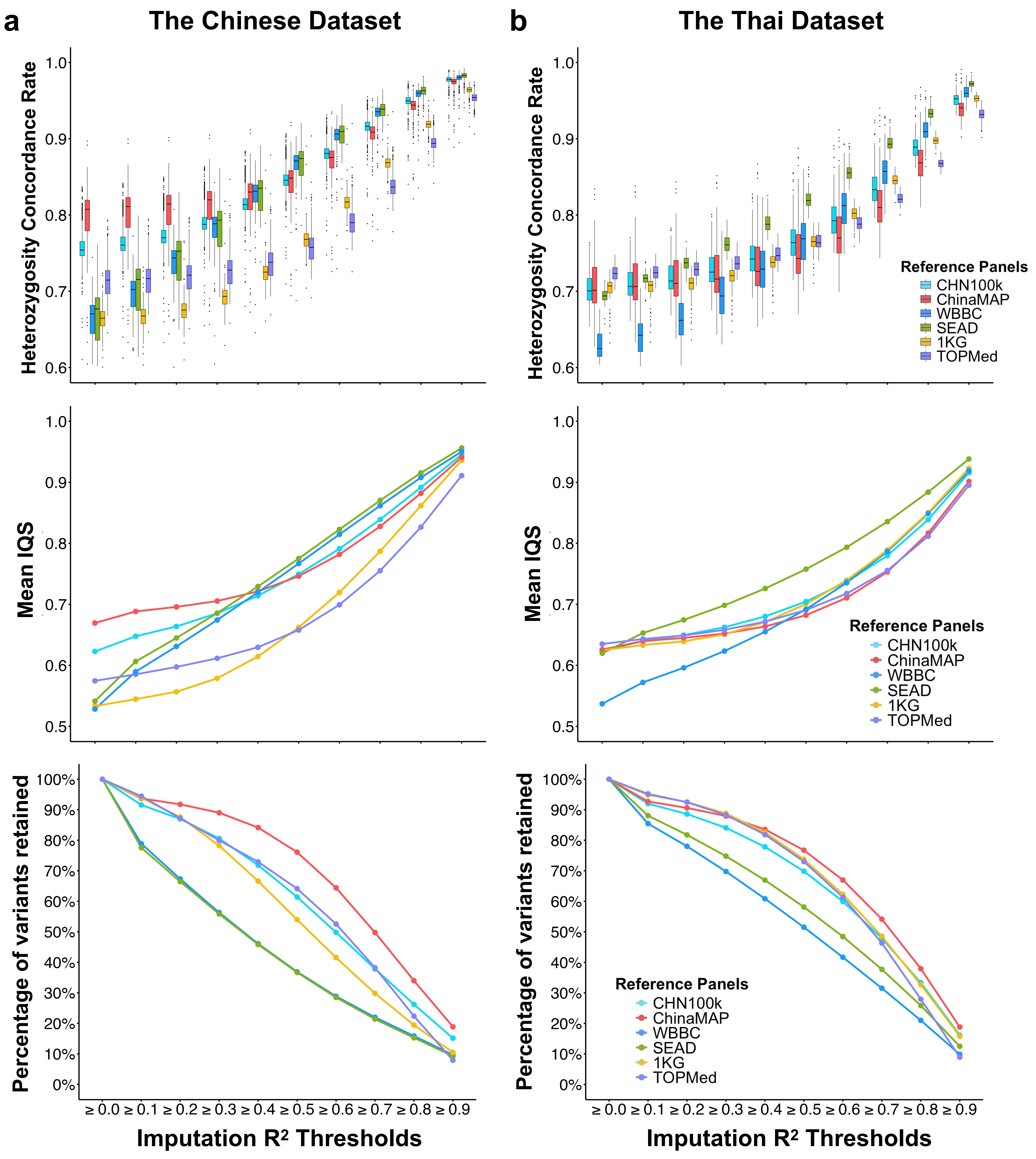


**Supplementary Figure 3 Evaluation of Imputation Quality for Low-Frequency Variants Across Reference Panels by Imputation $R^2$ Thresholds in Chinese (a) and Thai (b) WGS Datasets**

The X-axis represents imputation $R^2$ threshold categories, while the Y-axis in the top panel displays heterozygosity concordance rate for each sample. The middle panel shows the mean imputation quality score (IQS) and the bottom panel represents percentage of variants retained across $R^2$ thresholds. Low-frequency variants refer to variants with minor allele frequence less than 0.05.

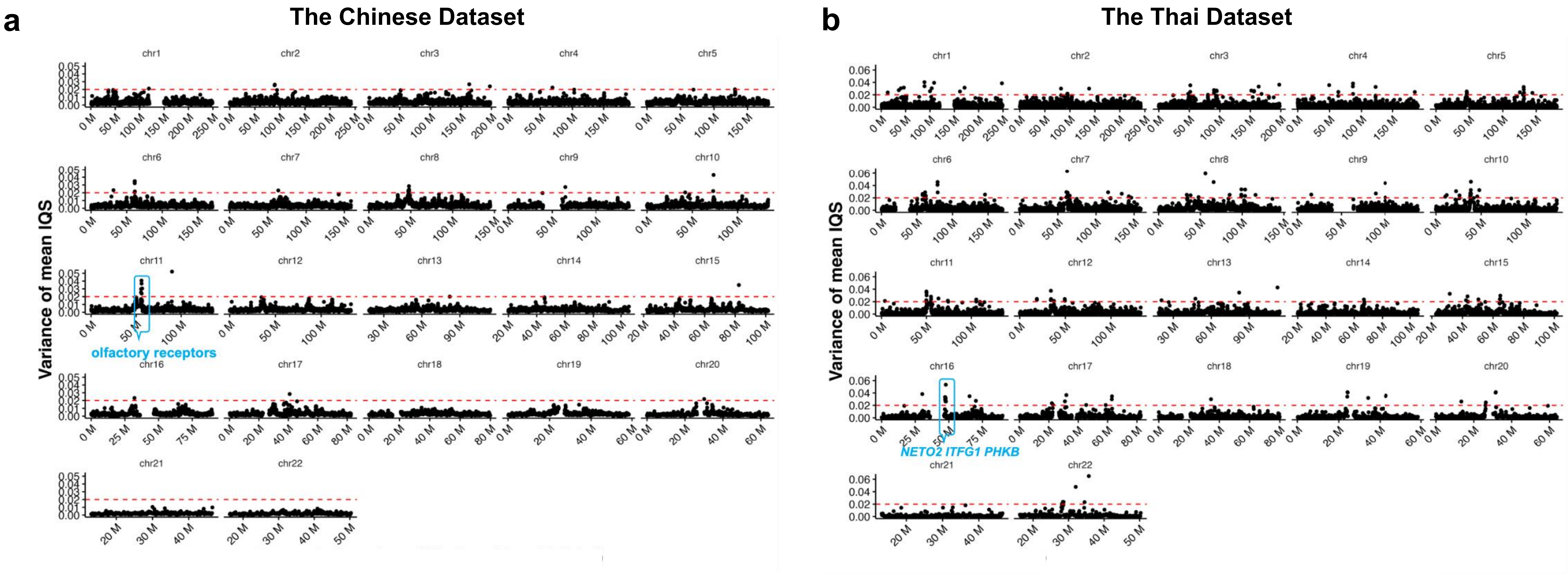


**Supplementary Figure 4 Overview of Imputation Variations in Reference Panels Across Genomic Regions in Chinese (a) and Thai (b) Datasets**

Each dot corresponds to a 100kb genomic region
