## Supplemental Tables for "Imputation Disparities Driven by Recent Selection and Their Impact on Disease Risk Estimation in East and Southeast Asian Populations"

**Supplementary Table 1 Summary of datasets used in this study**

| Platform | Chinese dataset | Thai dataset | Total |
|---|---|---|---|
| SNP array | 3,194 | 3,803 | 6,997 |
| WGS | 1,263 | 56 | 1,319 |
| Total individuals | 4,457 | 3,859 | 8,316 |

**Supplementary Table 2 Summary of imputation reference panels compared in our study**

| Reference Panels | Individuals | Percentage of Chinese | Percentage of East Asian | Average depth | Number of variants on autosomes |
|---|---|---|---|---|---|
| CHN100k | 25,734 | 100% | 100% | 40 | 111,918,691 |
| ChinaMAP | 10,155 | 100% | 100% | 40.8 | 59,010,860 |
| WBBC | 4,489 | 100% | 100% | 13.9 | 35,616,674 |
| SEAD | 11,067 | 68.6% | 82.3% | 13.6-36 | 80,367,720 |
| 1KG Phase3 v5 | 2,504 | 12.0% | 20.1% | 30 | 49,143,605 |
| TOPMed (r3) | 133,597 | 3.3% | 3.8% | 38 | 445,600,184 |

**Supplementary Table 3 Summary of imputed variant counts and their overlap with WGS data across reference panels**

| Dataset | CHN100k | ChinaMAP | WBBC | SEAD | 1KG | TOPMed |
|---|---|---|---|---|---|---|
| **The Chinese dataset** | | | | | | |
| Imputed variants | 111,107,029 | 58,666,899 | 35,284,726 | 80,020,879 | 48,545,446 | 421,738,285 |
| Variants overlapped with WGS | 29,324,080 | 25,186,782 | 18,890,460 | 22,494,639 | 13,625,262 | 25,396,871 |
| Variants overlapped with WGS (MAF ≤0.05) | 24,342,624 | 20,289,892 | 14,630,413 | 17,474,800 | 8,794,001 | 20,720,838 |
| Variants overlapped with WGS (MAF >0.05) | 4,981,456 | 4,896,890 | 4,260,047 | 5,019,839 | 4,831,261 | 4,676,033 |
| **The Thai dataset** | | | | | | |
| Imputed variants | 111,150,206 | 58,280,146 | 35,616,674 | 80,367,720 | 48,890,301 | 421,781,122 |
| Variants overlapped with WGS | 9,895,577 | 9,590,338 | 8,314,681 | 10,226,492 | 9,113,067 | 9,773,641 |
| Variants overlapped with WGS (MAF ≤0.05) | 4,783,181 | 4,490,612 | 3,774,391 | 4,811,739 | 3,934,446 | 4,770,982 |
| Variants overlapped with WGS (MAF >0.05) | 5,112,396 | 5,099,726 | 4,540,290 | 5,414,753 | 5,178,621 | 5,002,659 |

**Note:** Indels and SNVs within the human leukocyte antigens (HLA) region (chr6: 29,722,775-33,314,387) were excluded from the analysis. For the comparison, 42,750,853 SNVs outside the HLA region identified through WGS in the Chinese dataset and 12,597,713 SNVs outside the HLA region in the Thai dataset were included.

**Supplementary Table 4 Heterozygosity concordance rate across different $R^2$ thresholds**

| $R^2$ threshold | CHN100k | ChinaMAP | WBBC | SEAD | 1KG | TOPMed |
|---|---|---|---|---|---|---|
| **The Chinese dataset** | | | | | | |
| ⩾0.0 | 0.903 ± 0.010 | 0.918 ± 0.011 | 0.871 ± 0.011 | 0.880 ± 0.014 | 0.861 ± 0.005 | 0.873 ± 0.009 |
| ⩾0.1 | 0.904 ± 0.009 | 0.919 ± 0.011 | 0.874 ± 0.010 | 0.884 ± 0.014 | 0.862 ± 0.005 | 0.873 ± 0.009 |
| ⩾0.2 | 0.905 ± 0.009 | 0.919 ± 0.011 | 0.881 ± 0.010 | 0.891 ± 0.013 | 0.863 ± 0.005 | 0.874 ± 0.009 |
| ⩾0.3 | 0.908 ± 0.009 | 0.920 ± 0.011 | 0.892 ± 0.010 | 0.902 ± 0.012 | 0.866 ± 0.005 | 0.875 ± 0.009 |
| ⩾0.4 | 0.914 ± 0.009 | 0.922 ± 0.011 | 0.907 ± 0.009 | 0.916 ± 0.011 | 0.873 ± 0.005 | 0.878 ± 0.009 |
| ⩾0.5 | 0.924 ± 0.008 | 0.927 ± 0.010 | 0.924 ± 0.008 | 0.931 ± 0.009 | 0.885 ± 0.004 | 0.887 ± 0.008 |
| ⩾0.6 | 0.937 ± 0.007 | 0.936 ± 0.009 | 0.941 ± 0.007 | 0.947 ± 0.008 | 0.903 ± 0.004 | 0.903 ± 0.007 |
| ⩾0.7 | 0.952 ± 0.006 | 0.949 ± 0.008 | 0.957 ± 0.006 | 0.961 ± 0.006 | 0.924 ± 0.003 | 0.926 ± 0.006 |
| ⩾0.8 | 0.968 ± 0.005 | 0.965 ± 0.006 | 0.972 ± 0.004 | 0.975 ± 0.004 | 0.948 ± 0.003 | 0.953 ± 0.004 |
| ⩾0.9 | 0.985 ± 0.003 | 0.983 ± 0.004 | 0.986 ± 0.003 | 0.988 ± 0.002 | 0.974 ± 0.002 | 0.980 ± 0.002 |
| **The Thai dataset** | | | | | | |
| ⩾0.0 | 0.911 ± 0.012 | 0.913 ± 0.018 | 0.887 ± 0.014 | 0.911 ± 0.007 | 0.904 ± 0.005 | 0.907 ± 0.005 |
| ⩾0.1 | 0.911 ± 0.012 | 0.914 ± 0.018 | 0.891 ± 0.014 | 0.915 ± 0.007 | 0.904 ± 0.005 | 0.907 ± 0.005 |
| ⩾0.2 | 0.913 ± 0.012 | 0.915 ± 0.018 | 0.897 ± 0.014 | 0.919 ± 0.007 | 0.904 ± 0.005 | 0.908 ± 0.005 |
| ⩾0.3 | 0.916 ± 0.012 | 0.916 ± 0.017 | 0.905 ± 0.014 | 0.925 ± 0.007 | 0.906 ± 0.005 | 0.909 ± 0.005 |
| ⩾0.4 | 0.920 ± 0.012 | 0.918 ± 0.017 | 0.913 ± 0.013 | 0.932 ± 0.007 | 0.909 ± 0.005 | 0.912 ± 0.005 |
| ⩾0.5 | 0.927 ± 0.012 | 0.923 ± 0.017 | 0.924 ± 0.013 | 0.940 ± 0.007 | 0.916 ± 0.004 | 0.917 ± 0.005 |
| ⩾0.6 | 0.936 ± 0.011 | 0.930 ± 0.016 | 0.935 ± 0.012 | 0.949 ± 0.006 | 0.925 ± 0.004 | 0.925 ± 0.004 |
| ⩾0.7 | 0.947 ± 0.010 | 0.941 ± 0.015 | 0.948 ± 0.010 | 0.959 ± 0.005 | 0.937 ± 0.004 | 0.937 ± 0.004 |
| ⩾0.8 | 0.961 ± 0.008 | 0.955 ± 0.012 | 0.963 ± 0.008 | 0.971 ± 0.004 | 0.952 ± 0.004 | 0.955 ± 0.004 |
| ⩾0.9 | 0.979 ± 0.005 | 0.975 ± 0.007 | 0.980 ± 0.005 | 0.984 ± 0.003 | 0.972 ± 0.003 | 0.978 ± 0.003 |

**Note:** Mean ± stand deviation

**Supplementary Table 5 Mean IQS scores across different $R^2$ thresholds**

| Imputation $R^2$ | CHN100k | ChinaMAP | WBBC | SEAD | 1KG | TOPMed |
|---|---|---|---|---|---|---|
| **The Chinese dataset** | | | | | | |
| ⩾0.0 | 0.702 | 0.742 | 0.623 | 0.633 | 0.635 | 0.657 |
| ⩾0.1 | 0.720 | 0.756 | 0.669 | 0.682 | 0.646 | 0.666 |
| ⩾0.2 | 0.732 | 0.761 | 0.698 | 0.709 | 0.657 | 0.676 |
| ⩾0.3 | 0.747 | 0.768 | 0.728 | 0.738 | 0.675 | 0.688 |
| ⩾0.4 | 0.768 | 0.779 | 0.760 | 0.769 | 0.703 | 0.702 |
| ⩾0.5 | 0.794 | 0.797 | 0.795 | 0.803 | 0.740 | 0.724 |
| ⩾0.6 | 0.826 | 0.824 | 0.831 | 0.839 | 0.782 | 0.759 |
| ⩾0.7 | 0.863 | 0.859 | 0.870 | 0.877 | 0.830 | 0.805 |
| ⩾0.8 | 0.904 | 0.900 | 0.909 | 0.915 | 0.882 | 0.863 |
| ⩾0.9 | 0.948 | 0.947 | 0.949 | 0.953 | 0.938 | 0.929 |
| **The Thai dataset** | | | | | | |
| ⩾0.0 | 0.748 | 0.759 | 0.692 | 0.738 | 0.760 | 0.750 |
| ⩾0.1 | 0.758 | 0.768 | 0.713 | 0.756 | 0.766 | 0.755 |
| ⩾0.2 | 0.763 | 0.771 | 0.727 | 0.767 | 0.769 | 0.758 |
| ⩾0.3 | 0.771 | 0.775 | 0.744 | 0.781 | 0.775 | 0.763 |
| ⩾0.4 | 0.782 | 0.783 | 0.763 | 0.797 | 0.787 | 0.772 |
| ⩾0.5 | 0.798 | 0.795 | 0.785 | 0.815 | 0.803 | 0.784 |
| ⩾0.6 | 0.818 | 0.813 | 0.811 | 0.837 | 0.826 | 0.802 |
| ⩾0.7 | 0.845 | 0.839 | 0.842 | 0.864 | 0.854 | 0.828 |
| ⩾0.8 | 0.881 | 0.876 | 0.881 | 0.897 | 0.888 | 0.866 |
| ⩾0.9 | 0.931 | 0.926 | 0.931 | 0.941 | 0.932 | 0.923 |

**Supplementary Table 6 Heterozygosity concordance rate across different $R^2$ thresholds for low-frequency variants (MAF < 0.05)**

| Imputation $R^2$ | CHN100k | ChinaMAP | WBBC | SEAD | 1KG | TOPMed |
|---|---|---|---|---|---|---|
| **The Chinese dataset** | | | | | | |
| ⩾0.0 | 0.758 ± 0.021 | 0.802 ± 0.026 | 0.665 ± 0.024 | 0.669 ± 0.034 | 0.664 ± 0.013 | 0.715 ± 0.021 |
| ⩾0.1 | 0.765 ± 0.020 | 0.806 ± 0.025 | 0.698 ± 0.021 | 0.708 ± 0.032 | 0.667 ± 0.013 | 0.716 ± 0.021 |
| ⩾0.2 | 0.774 ± 0.020 | 0.810 ± 0.024 | 0.739 ± 0.020 | 0.745 ± 0.031 | 0.675 ± 0.013 | 0.721 ± 0.021 |
| ⩾0.3 | 0.791 ± 0.019 | 0.815 ± 0.023 | 0.785 ± 0.019 | 0.787 ± 0.028 | 0.692 ± 0.013 | 0.727 ± 0.021 |
| ⩾0.4 | 0.816 ± 0.018 | 0.826 ± 0.022 | 0.828 ± 0.017 | 0.829 ± 0.024 | 0.724 ± 0.013 | 0.738 ± 0.021 |
| ⩾0.5 | 0.847 ± 0.016 | 0.845 ± 0.020 | 0.869 ± 0.015 | 0.870 ± 0.019 | 0.767 ± 0.012 | 0.758 ± 0.020 |
| ⩾0.6 | 0.881 ± 0.014 | 0.874 ± 0.017 | 0.905 ± 0.013 | 0.907 ± 0.014 | 0.816 ± 0.010 | 0.791 ± 0.017 |
| ⩾0.7 | 0.917 ± 0.012 | 0.908 ± 0.014 | 0.934 ± 0.011 | 0.938 ± 0.010 | 0.868 ± 0.008 | 0.838 ± 0.014 |
| ⩾0.8 | 0.950 ± 0.009 | 0.943 ± 0.011 | 0.959 ± 0.008 | 0.963 ± 0.007 | 0.919 ± 0.006 | 0.895 ± 0.010 |
| ⩾0.9 | 0.977 ± 0.006 | 0.975 ± 0.007 | 0.980 ± 0.005 | 0.983 ± 0.004 | 0.964 ± 0.005 | 0.954 ± 0.006 |
| **The Thai dataset** | | | | | | |
| ⩾0.0 | 0.704 ± 0.026 | 0.711 ± 0.048 | 0.608 ± 0.032 | 0.694 ± 0.010 | 0.700 ± 0.020 | 0.724 ± 0.010 |
| ⩾0.1 | 0.711 ± 0.026 | 0.715 ± 0.048 | 0.637 ± 0.031 | 0.717 ± 0.010 | 0.701 ± 0.020 | 0.725 ± 0.010 |
| ⩾0.2 | 0.719 ± 0.027 | 0.720 ± 0.048 | 0.665 ± 0.034 | 0.737 ± 0.011 | 0.705 ± 0.021 | 0.729 ± 0.011 |
| ⩾0.3 | 0.730 ± 0.029 | 0.726 ± 0.048 | 0.697 ± 0.037 | 0.762 ± 0.012 | 0.715 ± 0.020 | 0.737 ± 0.011 |
| ⩾0.4 | 0.747 ± 0.032 | 0.737 ± 0.049 | 0.732 ± 0.039 | 0.790 ± 0.013 | 0.735 ± 0.019 | 0.749 ± 0.012 |
| ⩾0.5 | 0.770 ± 0.034 | 0.754 ± 0.051 | 0.771 ± 0.038 | 0.822 ± 0.014 | 0.763 ± 0.016 | 0.766 ± 0.011 |
| ⩾0.6 | 0.800 ± 0.034 | 0.780 ± 0.049 | 0.814 ± 0.035 | 0.857 ± 0.014 | 0.801 ± 0.012 | 0.789 ± 0.010 |
| ⩾0.7 | 0.840 ± 0.030 | 0.818 ± 0.045 | 0.860 ± 0.030 | 0.895 ± 0.011 | 0.846 ± 0.009 | 0.822 ± 0.008 |
| ⩾0.8 | 0.892 ± 0.022 | 0.874 ± 0.035 | 0.912 ± 0.022 | 0.934 ± 0.008 | 0.898 ± 0.006 | 0.868 ± 0.007 |
| ⩾0.9 | 0.953 ± 0.012 | 0.942 ± 0.018 | 0.961 ± 0.011 | 0.972 ± 0.005 | 0.953 ± 0.005 | 0.932 ± 0.009 |

**Note:** Mean ± stand deviation

**Supplementary Table 7 Mean IQS scores across different $R^2$ thresholds for low-frequency variants (MAF < 0.05)**

| Imputation $R^2$ | CHN100k | ChinaMAP | WBBC | SEAD | 1KG | TOPMed |
|---|---|---|---|---|---|---|
| **The Chinese dataset** | | | | | | |
| ≥0.0 | 0.623 | 0.669 | 0.528 | 0.541 | 0.533 | 0.575 |
| ≥0.1 | 0.648 | 0.688 | 0.590 | 0.606 | 0.545 | 0.585 |
| ≥0.2 | 0.664 | 0.696 | 0.631 | 0.645 | 0.557 | 0.597 |
| ≥0.3 | 0.685 | 0.705 | 0.674 | 0.686 | 0.579 | 0.611 |
| ≥0.4 | 0.714 | 0.721 | 0.720 | 0.729 | 0.614 | 0.630 |
| ≥0.5 | 0.749 | 0.746 | 0.767 | 0.775 | 0.662 | 0.658 |
| ≥0.6 | 0.791 | 0.782 | 0.815 | 0.823 | 0.719 | 0.699 |
| ≥0.7 | 0.839 | 0.828 | 0.862 | 0.870 | 0.787 | 0.755 |
| ≥0.8 | 0.892 | 0.882 | 0.908 | 0.915 | 0.862 | 0.827 |
| ≥0.9 | 0.945 | 0.941 | 0.950 | 0.956 | 0.936 | 0.911 |
| **The Thai dataset** | | | | | | |
| ≥0.0 | 0.623 | 0.626 | 0.537 | 0.620 | 0.625 | 0.635 |
| ≥0.1 | 0.640 | 0.640 | 0.572 | 0.653 | 0.633 | 0.643 |
| ≥0.2 | 0.649 | 0.645 | 0.596 | 0.674 | 0.639 | 0.649 |
| ≥0.3 | 0.662 | 0.652 | 0.623 | 0.698 | 0.651 | 0.658 |
| ≥0.4 | 0.680 | 0.664 | 0.655 | 0.726 | 0.671 | 0.672 |
| ≥0.5 | 0.704 | 0.682 | 0.692 | 0.758 | 0.700 | 0.691 |
| ≥0.6 | 0.737 | 0.711 | 0.735 | 0.794 | 0.740 | 0.718 |
| ≥0.7 | 0.780 | 0.753 | 0.787 | 0.836 | 0.789 | 0.755 |
| ≥0.8 | 0.839 | 0.817 | 0.849 | 0.884 | 0.850 | 0.812 |
| ≥0.9 | 0.916 | 0.901 | 0.919 | 0.938 | 0.923 | 0.895 |

**Supplementary Table 8 Summary of genomic regions with the best performance for different references**

| | CHN100k | ChinaMAP | WBBC | SEAD | 1KG | TOPMed | Total |
|---|---|---|---|---|---|---|---|
| Chinese | 1,486 | 24,343 | 62 | 25 | 283 | 153 | 26,352 |
| Thai | 3,289 | 8,239 | 63 | 3,031 | 8,098 | 3,309 | 26,029 |

**Note:** The imputation results for the Thai and Chinese datasets shared varying numbers of variants with the WGS data, leading to variations in the total number of windows between the two datasets.

**Supplementary Table 14 Imputed vs. Actual Genotype Probability Table for N Individuals**

| | | Actual | | | |
|---|---|---|---|---|---|
| | | 0/0 | 0/1 | 1/1 | Total |
| Imputed | 0/0 | $\sum_{n=1}^{N_{11}} p_{11\_n}$ | $\sum_{n=1}^{N_{12}} p_{12\_n}$ | $\sum_{n=1}^{N_{13}} p_{13\_n}$ | $\sum_{j=1}^{3}\sum_{n=1}^{N_{1j}} p_{1j\_n} = Y_1$ |
| | 0/1 | $\sum_{n=1}^{N_{21}} p_{21\_n}$ | $\sum_{n=1}^{N_{22}} p_{22\_n}$ | $\sum_{n=1}^{N_{23}} p_{23\_n}$ | $\sum_{j=1}^{3}\sum_{n=1}^{N_{2j}} p_{2j\_n} = Y_2$ |
| | 1/1 | $\sum_{n=1}^{N_{31}} p_{31\_n}$ | $\sum_{n=1}^{N_{32}} p_{32\_n}$ | $\sum_{n=1}^{N_{33}} p_{33\_n}$ | $\sum_{j=1}^{3}\sum_{n=1}^{N_{3j}} p_{3j\_n} = Y_3$ |
| | Total | $\sum_{i=1}^{3}\sum_{n=1}^{N_{i1}} p_{i1\_n} = W_1$ | $\sum_{i=1}^{3}\sum_{n=1}^{N_{i2}} p_{i2\_n} = W_2$ | $\sum_{i=1}^{3}\sum_{n=1}^{N_{i3}} p_{i3\_n} = W_3$ | $N$ |

**Note:** $p_{ij\_n}$ denotes the genotype probability that the n$^{th}$ individual is imputed with genotype $i$ while actually possessing genotype $j$, where 1 corresponds to the homozygous reference genotype (0/0), 2 refers to the heterozygous genotype (0/1), and 3 denotes the alternative homozygous genotype (1/1). Each cell within this matrix represents the cumulative genotype probabilities for each combination of actual and imputed genotypic classes.

**Supplementary Table 15 Summary of SLE-associated variants used in this study**

| SNP | Alt Allele | Effect Allele | OR | P | BETA | PMID |
|---|---|---|---|---|---|---|
| rs10018951 | C | T | 1.310 | 1.18E-14 | 0.270 | *PMID:33536424* |
| rs10239000 | A | G | 0.841 | 1.51E-25 | -0.173 | *PMID:33536424* |
| rs10807150 | C | T | 0.800 | 6.06E-16 | -0.223 | *PMID:33536424* |
| rs10823829 | T | C | 1.099 | 1.05E-09 | 0.094 | *PMID:33536424* |
| rs10896045 | A | G | 0.856 | 6.59E-26 | -0.155 | *PMID:33536424* |
| rs10995261 | C | T | 0.909 | 2.57E-08 | -0.095 | *PMID:33536424* |
| rs11032994 | G | A | 1.147 | 1.12E-17 | 0.137 | *PMID:33536424* |
| rs11673604 | T | C | 0.874 | 4.21E-12 | -0.135 | *PMID:33536424* |
| rs116991837 | G | A | 2.654 | 3.82E-15 | 0.976 | *PMID:33536424* |
| rs117821148 | C | T | 1.460 | 4.80E-08 | 0.378 | *PMID:33536424* |
| rs118075465 | G | A | 1.140 | 1.16E-10 | 0.131 | *PMID:33536424* |
| rs12120358 | A | T | 1.188 | 2.96E-11 | 0.172 | *PMID:33536424* |
| rs12153670 | A | G | 1.150 | 7.65E-10 | 0.140 | *PMID:33536424* |
| rs1234314 | C | G | 1.389 | 5.84E-11 | 0.329 | *PMID:33536424* |
| rs12461589 | C | T | 0.898 | 5.00E-10 | -0.108 | *PMID:33536424* |
| rs12822507 | A | G | 0.860 | 2.20E-08 | -0.151 | *PMID:33536424* |
| rs12900339 | A | G | 0.850 | 4.73E-10 | -0.163 | *PMID:33536424* |
| rs12900640 | A | C | 0.908 | 2.42E-11 | -0.096 | *PMID:33536424* |
| rs13116227 | C | T | 1.340 | 3.05E-11 | 0.293 | *PMID:33536424* |
| rs13244581 | G | C | 0.667 | 1.59E-20 | -0.405 | *PMID:33536424* |
| rs13259960 | A | G | 1.350 | 1.03E-11 | 0.300 | *PMID:33536424* |
| rs13306575 | G | A | 1.311 | 2.28E-14 | 0.271 | *PMID:33536424* |

| | | | | | | |
|---|---|---|---|---|---|---|
| rs138305363 | A | G | 1.608 | 4.07E-08 | 0.475 | *PMID:33536424* |
| rs143176121 | T | C | 3.425 | 1.58E-66 | 1.231 | *PMID:33536424* |
| rs145720245 | G | A | 4.000 | 2.80E-10 | 1.386 | *PMID:33536424* |
| rs146063533 | C | T | 1.612 | 9.44E-16 | 0.477 | *PMID:33536424* |
| rs150724213 | G | A | 3.882 | 2.51E-15 | 1.356 | *PMID:33536424* |
| rs16902895 | A | G | 0.891 | 1.48E-13 | -0.115 | *PMID:33536424* |
| rs17374162 | G | A | 0.917 | 3.02E-09 | -0.087 | *PMID:33536424* |
| rs1887428 | G | C | 0.806 | 4.49E-14 | -0.215 | *PMID:33536424* |
| rs2039982 | C | T | 1.242 | 1.85E-37 | 0.217 | *PMID:33536424* |
| rs218174 | A | G | 0.892 | 1.83E-13 | -0.114 | *PMID:33536424* |
| rs2233302 | C | G | 0.760 | 1.17E-08 | -0.274 | *PMID:33536424* |
| rs2238577 | C | T | 0.885 | 1.83E-14 | -0.122 | *PMID:33536424* |
| rs2272736 | G | A | 0.819 | 6.37E-11 | -0.200 | *PMID:33536424* |
| rs231694 | T | C | 0.900 | 9.71E-09 | -0.105 | *PMID:33536424* |
| rs2362475 | A | C | 1.176 | 2.00E-09 | 0.163 | *PMID:33536424* |
| rs2421184 | A | G | 0.840 | 4.67E-12 | -0.174 | *PMID:33536424* |
| rs244689 | A | G | 0.775 | 1.21E-20 | -0.255 | *PMID:33536424* |
| rs2540119 | T | C | 0.921 | 3.51E-08 | -0.083 | *PMID:33536424* |
| rs2549002 | C | A | 0.905 | 2.40E-10 | -0.100 | *PMID:33536424* |
| rs2671655 | C | T | 1.087 | 4.60E-08 | 0.083 | *PMID:33536424* |
| rs2714333 | T | C | 0.322 | 1.00E-08 | -1.135 | *PMID:33536424* |
| rs2819426 | G | C | 0.824 | 2.51E-30 | -0.194 | *PMID:33536424* |
| rs28364822 | T | A | 1.672 | 1.19E-17 | 0.514 | *PMID:33536424* |
| rs2855772 | T | C | 1.400 | 1.21E-15 | 0.336 | *PMID:33536424* |

| | | | | | | |
|---|---|---|---|---|---|---|
| rs34330 | T | C | 1.190 | 4.80E-12 | 0.174 | |
| rs35032408 | T | G | 0.690 | 2.84E-08 | -0.371 | *PMID:33536424* |
| rs35966917 | A | G | 1.094 | 4.66E-09 | 0.090 | *PMID:33536424* |
| rs35985016 | A | G | 1.186 | 1.95E-08 | 0.171 | *PMID:33536424* |
| rs372605131 | T | C | 1.780 | 1.20E-08 | 0.577 | *PMID:33536424* |
| rs3748079 | C | T | 0.532 | 1.78E-08 | -0.631 | *PMID:33536424* |
| rs3750996 | A | G | 0.857 | 1.89E-12 | -0.154 | *PMID:33536424* |
| rs3794986 | G | T | 0.890 | 1.46E-14 | -0.117 | *PMID:33536424* |
| rs3806357 | G | A | 1.106 | 4.25E-09 | 0.101 | *PMID:33536424* |
| rs3857496 | T | C | 0.896 | 7.66E-09 | -0.110 | *PMID:33536424* |
| rs3999421 | A | T | 1.099 | 1.29E-09 | 0.094 | *PMID:33536424* |
| rs41298401 | C | G | 0.773 | 5.04E-45 | -0.258 | *PMID:33536424* |
| rs4251697 | G | A | 0.637 | 1.17E-43 | -0.451 | *PMID:33536424* |
| rs447632 | A | G | 1.175 | 7.23E-28 | 0.161 | *PMID:33536424* |
| rs4573208 | G | A | 1.173 | 1.53E-15 | 0.160 | *PMID:33536424* |
| rs4598207 | A | T | 0.750 | 4.12E-60 | -0.287 | *PMID:33536424* |
| rs4622329 | G | A | 0.894 | 4.00E-15 | -0.112 | *PMID:33536424* |
| rs4643809 | C | T | 0.846 | 3.53E-24 | -0.167 | *PMID:33536424* |
| rs4649203 | G | A | 0.862 | 9.90E-09 | -0.148 | *PMID:33536424* |
| rs4801882 | G | A | 0.882 | 1.86E-18 | -0.126 | *PMID:33536424* |
| rs4819670 | T | C | 0.869 | 5.53E-11 | -0.141 | *PMID:33536424* |
| rs4821116 | C | T | 1.240 | 8.86E-46 | 0.215 | *PMID:33536424* |
| rs4852324 | T | C | 0.790 | 5.70E-14 | -0.236 | *PMID:33536424* |
| rs4930642 | A | G | 0.873 | 6.16E-13 | -0.135 | *PMID:33536424* |

| | | | | | | |
|---|---|---|---|---|---|---|
| rs4936441 | C | G | 1.215 | 5.71E-16 | 0.195 | *PMID:33536424* |
| rs529561493 | C | A | 3.660 | 9.80E-09 | 1.297 | *PMID:33536424* |
| rs530634980 | C | T | 2.016 | 5.10E-18 | 0.701 | *PMID:33536424* |
| rs55882956 | G | A | 0.674 | 1.23E-16 | -0.395 | *PMID:33536424* |
| rs57095329 | A | G | 1.290 | 2.74E-08 | 0.255 | *PMID:33536424* |
| rs57141708 | G | A | 1.183 | 6.84E-22 | 0.168 | *PMID:33536424* |
| rs58107865 | G | C | 0.802 | 6.57E-25 | -0.221 | *PMID:33536424* |
| rs58164562 | T | C | 1.121 | 3.14E-12 | 0.114 | *PMID:33536424* |
| rs5826945 | A | T | 1.196 | 9.67E-11 | 0.179 | *PMID:33536424* |
| rs61759532 | C | T | 1.235 | 2.79E-11 | 0.211 | *PMID:33536424* |
| rs6533951 | A | G | 0.900 | 1.25E-10 | -0.105 | *PMID:33536424* |
| rs6539078 | T | C | 1.119 | 9.49E-14 | 0.112 | *PMID:33536424* |
| rs684150 | C | T | 0.914 | 4.32E-10 | -0.090 | *PMID:33536424* |
| rs6841907 | T | C | 1.104 | 1.10E-09 | 0.099 | *PMID:33536424* |
| rs707149 | A | G | 1.235 | 3.58E-08 | 0.211 | *PMID:33536424* |
| rs7072606 | T | C | 1.131 | 2.22E-12 | 0.123 | *PMID:33536424* |
| rs7170151 | C | T | 1.107 | 3.20E-12 | 0.102 | *PMID:33536424* |
| rs73954925 | C | G | 0.855 | 5.11E-11 | -0.156 | *PMID:33536424* |
| rs74989671 | A | G | 1.545 | 1.61E-08 | 0.435 | *PMID:33536424* |
| rs75362385 | G | T | 0.887 | 8.40E-13 | -0.120 | *PMID:33536424* |
| rs7565158 | G | T | 1.096 | 2.88E-10 | 0.092 | *PMID:33536424* |
| rs7572733 | C | T | 1.143 | 1.25E-14 | 0.134 | *PMID:33536424* |
| rs75773410 | A | G | 1.292 | 3.84E-15 | 0.256 | *PMID:33536424* |
| rs76107698 | G | C | 0.788 | 1.85E-30 | -0.238 | *PMID:33536424* |

| | | | | | | |
|---|---|---|---|---|---|---|
| rs7637844 | A | C | 1.140 | 1.28E-08 | 0.131 | *PMID:33536424* |
| rs77009341 | G | C | 2.089 | 6.39E-62 | 0.737 | *PMID:33536424* |
| rs7725218 | G | A | 1.132 | 2.47E-17 | 0.124 | *PMID:33536424* |
| rs77285596 | T | G | 0.734 | 2.20E-19 | -0.309 | *PMID:33536424* |
| rs77448389 | A | G | 1.170 | 7.30E-10 | 0.157 | *PMID:33536424* |
| rs77465633 | C | A | 1.340 | 6.99E-18 | 0.293 | *PMID:33536424* |
| rs77885959 | T | G | 0.590 | 3.16E-17 | -0.527 | *PMID:33536424* |
| rs77971648 | T | C | 0.775 | 3.16E-23 | -0.255 | *PMID:33536424* |
| rs780669 | C | T | 1.160 | 4.83E-09 | 0.148 | *PMID:33536424* |
| rs7858766 | T | C | 0.878 | 2.25E-15 | -0.130 | *PMID:33536424* |
| rs7902146 | C | T | 0.900 | 3.34E-12 | -0.105 | *PMID:33536424* |
| rs7911501 | G | A | 2.100 | 2.00E-09 | 0.742 | *PMID:33536424* |
| rs79171842 | A | T | 2.469 | 3.02E-23 | 0.904 | *PMID:33536424* |
| rs79401250 | T | G | 0.853 | 1.48E-12 | -0.159 | *PMID:33536424* |
| rs794368 | A | G | 0.844 | 1.54E-26 | -0.170 | *PMID:33536424* |
| rs79774308 | A | G | 0.689 | 5.76E-10 | -0.373 | *PMID:33536424* |
| rs8016947 | T | G | 1.205 | 1.08E-13 | 0.186 | *PMID:33536424* |
| rs9295676 | G | T | 1.105 | 7.94E-11 | 0.100 | *PMID:33536424* |
| rs9322454 | G | A | 1.090 | 2.42E-08 | 0.086 | *PMID:33536424* |
| rs933717 | T | C | 7.692 | 2.36E-10 | 2.040 | *PMID:33536424* |
| rs9488914 | C | T | 0.862 | 1.14E-08 | -0.149 | *PMID:33536424* |
| rs9503037 | A | G | 1.135 | 1.36E-15 | 0.127 | *PMID:33536424* |
| rs956237 | G | A | 1.107 | 4.47E-11 | 0.102 | *PMID:33536424* |
| rs9651076 | A | G | 0.895 | 3.26E-13 | -0.111 | *PMID:33536424* |

| | | | | | | |
|---|---|---|---|---|---|---|
| rs9736939 | G | A | 1.265 | 1.23E-58 | 0.235 | *PMID:33536424* |
| rs117026326 | C | T | 2.994 | 7.35E-27 | 1.097 | *PMID:39624492* |
| rs1143679 | G | A | 2.276 | 2.36E-06 | 0.822 | *PMID:39624492* |
| rs2230926 | T | G | 2.001 | 5.76E-46 | 0.694 | *PMID:39624492* |
| rs4728142 | G | A | 1.562 | 2.55E-45 | 0.446 | *PMID:39624492* |
| rs11889341 | C | T | 1.543 | 3.79E-79 | 0.434 | *PMID:39624492* |
| rs2205960 | G | T | 1.422 | 5.82E-46 | 0.352 | *PMID:39624492* |
| rs2736340 | C | T | 1.403 | 1.29E-38 | 0.339 | *PMID:39624492* |
| rs73135369 | T | C | 1.400 | 2.15E-11 | 0.336 | *PMID:39624492* |
| rs1128334 | C | T | 1.373 | 1.70E-42 | 0.317 | *PMID:39624492* |
| rs3734266 | T | C | 1.339 | 1.96E-18 | 0.292 | *PMID:39624492* |
| rs2280381 | C | T | 1.323 | 6.75E-15 | 0.280 | *PMID:39624492* |
| rs7726414 | C | T | 1.302 | 2.26E-10 | 0.264 | *PMID:39624492* |
| rs1385374 | C | T | 1.290 | 1.35E-20 | 0.255 | *PMID:39624492* |
| rs1418190 | C | T | 1.268 | 1.13E-15 | 0.237 | *PMID:39624492* |
| rs10036748 | C | T | 1.250 | 3.22E-16 | 0.223 | *PMID:39624492* |
| rs11773745 | A | G | 1.239 | 2.52E-11 | 0.214 | *PMID:39624492* |
| rs6927090 | G | T | 1.233 | 3.48E-10 | 0.209 | *PMID:39624492* |
| rs35426045 | G | A | 1.224 | 5.21E-07 | 0.202 | *PMID:39624492* |
| rs7444 | T | C | 1.215 | 8.56E-18 | 0.195 | *PMID:39624492* |
| rs2841280 | G | C | 1.210 | 6.12E-11 | 0.191 | *PMID:39624492* |
| rs7329174 | A | G | 1.192 | 1.17E-07 | 0.176 | *PMID:39624492* |
| rs9782955 | T | C | 1.184 | 2.60E-04 | 0.169 | *PMID:39624492* |
| rs2297550 | C | G | 1.179 | 1.57E-08 | 0.165 | *PMID:39624492* |

| | | | | | | |
|---|---|---|---|---|---|---|
| rs2381401 | C | T | 1.179 | 1.11E-07 | 0.165 | *PMID:39624492* |
| rs34562254 | G | A | 1.176 | 2.88E-08 | 0.162 | *PMID:39624492* |
| rs1547624 | A | T | 1.172 | 4.55E-08 | 0.159 | *PMID:39624492* |
| rs12132445 | G | A | 1.171 | 6.75E-08 | 0.158 | *PMID:39624492* |
| rs61616683 | C | T | 1.168 | 8.15E-08 | 0.155 | *PMID:39624492* |
| rs10999979 | C | A | 1.163 | 1.01E-07 | 0.151 | *PMID:39624492* |
| rs7579944 | T | C | 1.163 | 3.17E-11 | 0.151 | *PMID:39624492* |
| rs2732552 | T | C | 1.162 | 1.74E-08 | 0.150 | *PMID:39624492* |
| rs7726159 | C | A | 1.156 | 4.72E-10 | 0.145 | *PMID:39624492* |
| rs2934498 | A | G | 1.151 | 4.55E-06 | 0.141 | *PMID:39624492* |
| rs76725306 | G | A | 1.150 | 1.38E-06 | 0.140 | *PMID:39624492* |
| rs11679484 | C | A | 1.147 | 5.59E-05 | 0.137 | *PMID:39624492* |
| rs4639966 | T | C | 1.147 | 5.20E-09 | 0.137 | *PMID:39624492* |
| rs4948496 | T | C | 1.143 | 6.00E-06 | 0.134 | *PMID:39624492* |
| rs8035957 | T | C | 1.136 | 2.43E-08 | 0.128 | *PMID:39624492* |
| rs7941765 | T | C | 1.130 | 4.20E-04 | 0.122 | *PMID:39624492* |
| rs1801274 | A | G | 1.127 | 1.79E-05 | 0.120 | *PMID:39624492* |
| rs438613 | T | C | 1.125 | 6.46E-08 | 0.118 | *PMID:39624492* |
| rs930297 | C | T | 1.120 | 2.67E-02 | 0.113 | *PMID:39624492* |
| rs6074813 | G | T | 1.118 | 8.49E-05 | 0.112 | *PMID:39624492* |
| rs4978037 | T | C | 1.117 | 7.36E-05 | 0.111 | *PMID:39624492* |
| rs494003 | G | A | 1.113 | 2.08E-02 | 0.107 | *PMID:39624492* |
| rs1405209 | T | C | 1.106 | 3.10E-04 | 0.101 | *PMID:39624492* |
| rs2322659 | T | C | 1.106 | 7.38E-06 | 0.101 | *PMID:39624492* |

| | | | | | | |
|---|---|---|---|---|---|---|
| rs6762714 | C | T | 1.106 | 9.41E-03 | 0.101 | *PMID:39624492* |
| rs2384991 | A | C | 1.101 | 2.88E-05 | 0.096 | *PMID:39624492* |
| rs597325 | A | G | 1.094 | 8.56E-05 | 0.090 | *PMID:39624492* |
| rs4690229 | A | T | 1.090 | 4.14E-03 | 0.086 | *PMID:39624492* |
| rs1990760 | C | T | 1.078 | 6.94E-03 | 0.075 | *PMID:39624492* |
| rs2289583 | C | A | 1.068 | 6.10E-02 | 0.066 | *PMID:39624492* |
| rs6740462 | C | A | 1.066 | 3.38E-02 | 0.064 | *PMID:39624492* |
| rs3024505 | G | A | 1.021 | 7.41E-01 | 0.021 | *PMID:39624492* |
| rs2428 | C | T | 1.017 | 6.92E-01 | 0.017 | *PMID:39624492* |
| rs2304256 | C | A | 0.993 | 7.58E-01 | -0.007 | *PMID:39624492* |
| rs2366293 | G | C | 0.970 | 7.23E-01 | -0.030 | *PMID:39624492* |
| rs9899849 | G | A | 0.969 | 3.32E-01 | -0.031 | *PMID:39624492* |
| rs4902562 | A | G | 0.967 | 2.70E-01 | -0.034 | *PMID:39624492* |
| rs17321999 | C | A | 0.939 | 1.24E-01 | -0.063 | *PMID:39624492* |
| rs6445975 | G | T | 0.933 | 9.66E-03 | -0.069 | *PMID:39624492* |
| rs12148050 | A | G | 0.930 | 1.04E-03 | -0.073 | *PMID:39624492* |
| rs4810485 | T | G | 0.929 | 1.20E-03 | -0.074 | *PMID:39624492* |
| rs13238909 | G | A | 0.915 | 1.97E-01 | -0.089 | *PMID:39624492* |
| rs3768792 | G | A | 0.914 | 9.61E-03 | -0.090 | *PMID:39624492* |
| rs4690055 | G | A | 0.911 | 4.14E-05 | -0.093 | *PMID:39624492* |
| rs2305772 | G | A | 0.905 | 1.73E-05 | -0.100 | *PMID:39624492* |
| rs10750836 | C | T | 0.897 | 1.67E-04 | -0.109 | *PMID:39624492* |
| rs10936599 | C | T | 0.895 | 4.50E-07 | -0.111 | *PMID:39624492* |
| rs1170426 | C | T | 0.893 | 1.44E-03 | -0.113 | *PMID:39624492* |

| | | | | | | |
|---|---|---|---|---|---|---|
| rs763361 | T | C | 0.893 | 9.76E-07 | -0.113 | *PMID:39624492* |
| rs13260060 | G | A | 0.889 | 1.92E-08 | -0.118 | *PMID:39624492* |
| rs11603023 | T | C | 0.886 | 1.27E-06 | -0.121 | *PMID:39624492* |
| rs2445610 | A | G | 0.886 | 5.26E-08 | -0.121 | *PMID:39624492* |
| rs10419308 | G | A | 0.885 | 3.60E-02 | -0.122 | *PMID:39624492* |
| rs6702599 | A | C | 0.882 | 3.56E-02 | -0.126 | *PMID:39624492* |
| rs2731783 | A | G | 0.881 | 2.42E-05 | -0.127 | *PMID:39624492* |
| rs3087243 | G | A | 0.879 | 6.44E-06 | -0.129 | *PMID:39624492* |
| rs3795310 | C | T | 0.877 | 1.40E-03 | -0.131 | *PMID:39624492* |
| rs4745876 | G | A | 0.876 | 1.36E-06 | -0.132 | *PMID:39624492* |
| rs6871748 | T | C | 0.875 | 1.44E-05 | -0.134 | *PMID:39624492* |
| rs405858 | C | T | 0.872 | 1.70E-08 | -0.137 | *PMID:39624492* |
| rs1016140 | G | T | 0.870 | 2.95E-10 | -0.139 | *PMID:39624492* |
| rs1885889 | A | G | 0.870 | 7.93E-10 | -0.139 | *PMID:39624492* |
| rs223881 | T | C | 0.869 | 6.97E-10 | -0.140 | *PMID:39624492* |
| rs7975703 | C | T | 0.869 | 1.47E-06 | -0.140 | *PMID:39624492* |
| rs3760667 | C | T | 0.863 | 1.38E-06 | -0.147 | *PMID:39624492* |
| rs7815944 | A | G | 0.862 | 1.78E-09 | -0.149 | *PMID:39624492* |
| rs13344313 | G | A | 0.860 | 1.35E-05 | -0.151 | *PMID:39624492* |
| rs2009453 | C | T | 0.859 | 1.96E-11 | -0.152 | *PMID:39624492* |
| rs28411034 | G | A | 0.859 | 5.80E-06 | -0.152 | *PMID:39624492* |
| rs564799 | C | T | 0.859 | 8.84E-06 | -0.152 | *PMID:39624492* |
| rs869310 | T | G | 0.856 | 1.11E-05 | -0.155 | *PMID:39624492* |
| rs10028805 | G | A | 0.854 | 8.41E-11 | -0.158 | *PMID:39624492* |

| | | | | | | |
|---|---|---|---|---|---|---|
| rs849142 | T | C | 0.854 | 2.68E-01 | -0.158 | *PMID:39624492* |
| rs4592664 | T | C | 0.852 | 2.37E-06 | -0.160 | *PMID:39624492* |
| rs6705628 | C | T | 0.851 | 4.71E-08 | -0.161 | *PMID:39624492* |
| rs9630991 | G | A | 0.850 | 3.34E-06 | -0.163 | *PMID:39624492* |
| rs4697651 | C | T | 0.848 | 4.49E-05 | -0.165 | *PMID:39624492* |
| rs34889541 | G | A | 0.847 | 1.38E-04 | -0.166 | *PMID:39624492* |
| rs12093154 | G | A | 0.844 | 4.12E-06 | -0.170 | *PMID:39624492* |
| rs12599402 | T | C | 0.843 | 1.63E-13 | -0.171 | *PMID:39624492* |
| rs463426 | T | C | 0.841 | 3.60E-14 | -0.173 | *PMID:39624492* |
| rs10845606 | C | A | 0.829 | 3.80E-09 | -0.188 | *PMID:39624492* |
| rs1131265 | G | C | 0.827 | 1.37E-14 | -0.190 | *PMID:39624492* |
| rs11644034 | G | A | 0.822 | 2.33E-04 | -0.196 | *PMID:39624492* |
| rs12802200 | C | A | 0.818 | 1.23E-02 | -0.201 | *PMID:39624492* |
| rs1308020 | G | A | 0.818 | 1.83E-09 | -0.201 | *PMID:39624492* |
| rs548234 | C | T | 0.801 | 2.39E-20 | -0.222 | *PMID:39624492* |
| rs1061502 | T | C | 0.795 | 2.25E-02 | -0.229 | *PMID:39624492* |
| rs7097397 | G | A | 0.783 | 4.79E-25 | -0.245 | *PMID:39624492* |
| rs2431697 | T | C | 0.779 | 1.57E-13 | -0.250 | *PMID:39624492* |
| rs11235604 | C | T | 0.778 | 1.26E-06 | -0.251 | *PMID:39624492* |
| rs729302 | A | C | 0.763 | 1.74E-28 | -0.270 | *PMID:39624492* |
| rs549669428 | T | G | 0.754 | 1.85E-05 | -0.282 | *PMID:39624492* |
| rs4917014 | T | G | 0.753 | 5.18E-29 | -0.284 | *PMID:39624492* |
| rs11264750 | A | G | 0.751 | 1.95E-12 | -0.286 | *PMID:39624492* |
| rs9387400 | C | A | 0.743 | 3.14E-08 | -0.297 | *PMID:39624492* |

| | | | | | | |
|---|---|---|---|---|---|---|
| rs13385731 | T | C | 0.710 | 2.96E-23 | -0.342 | *PMID:39624492* |
